# Self-supervised AI reveals a lethal discohesive phenotype in lung adenocarcinoma

**DOI:** 10.1101/2025.11.12.688049

**Authors:** Kai Rakovic, Alexandrina Pancheva, Adalberto Claudio Quiros, Jose Coelho-Lima, Zhangyi He, David A Dorward, Marco Sereno, Claire Wilson, Craig Dick, David A Moore, Fiona Ballantyne, Catherine Ficken, Ana Teodósio, Silvia Martinelli, Rachel Baird, Leah Officer-Jones, Ian R Powley, David Chang, Crispin J Miller, Ke Yuan, John Le Quesne

## Abstract

Applications of artificial intelligence (AI) to histopathology are now common, but most require supervision which inherently limits their scope. By using self-supervised learning (SSL), we discover and quantify the full range of histopathological appearances in a disease, and associate them with clinicopathological ground truths such as prognosis. We used this approach to discover under-appreciated morphologies of lung adenocarcinoma (LUAD), using a highly characterised resected tumour cohort of over 4000 slides from over 1000 patients. By constructing an authoritative lexicon of recurrent LUAD appearances, we *ab initio* discovered several stromal morphologies strongly predictive of outcome. With multimodal data integration and external dataset validation, we propose that epithelial discohesion is lethal, but only in the context of immunologically cold stroma. Both these morphological features are independent of current prognostic schema. Crucially, we describe these features in the context of real-world diagnostic histopathology, giving them immediate clinical translatability.

**Statement of Significance:** Histopathology relies heavily on epithelial morphology, often neglecting the stroma. We use self-supervised AI to identify under-appreciated morphologies in LUAD linked to poor outcome and validate these observations in an external cohort, demonstrating the utility of self-supervised AI as a powerful biological discovery tool.

## Introduction

The prognosis of patients with lung cancer remains dismal despite advances in therapy, and it is still the leading cause of cancer-related mortality worldwide [1]. Within common types of lung cancer, only non-small cell carcinomas are amenable to surgical cure, and the most common histological subtype is lung adenocarcinoma (LUAD). There is an ever-expanding battery of personalised therapeutic options in LUAD which are starting to improve the outlook [2], but these all rely on expensive and time-consuming molecular tests to subclassify patients rather than tissue morphology. The histopathological classification of LUAD is based on the epithelial growth pattern, and this system is well-established as a prognosticator in LUAD [3]. Although gradual refinements have been made over time, most recently with a new grading system from the International Association for the Study of Lung Cancer (IASLC) [4], the underlying morphological definitions of grade have not been substantially updated for a decade [5], however there have been recent refinements in difficult tasks such as defining invasion in early lesions [6]. This underlines the fact that there is a wealth of morphological information still locked in cheap and readily accessible haematoxylin and eosin-stained (H&E) slides, only a fraction of which is currently accessible for patient diagnosis, prognosis and treatment.

Artificial intelligence (AI) in histopathology is typically supervised, with a pre-defined task that the AI must be optimised for. Supervised learning is inherently limited by existing knowledge. In other words, the models are trained to replicate human performance rather than exceed it. Self-supervised learning (SSL) allows us to train AI models which do not require up-front hypotheses or annotations. While SSL is now extensively utilised in computational pathology with the advent of foundation models [7–10] these are generalist models optimised for common benchmarking tasks across multiple cancer types which may paradoxically limit their generalisability [11]. While they are undoubtedly powerful resources, these models are very broad, making them less suited to the discovery of granular morphological patterns across single cancer types. Furthermore, they lack interpretability which hinders their utility, especially for biological discovery.

While there is strong evidence of the importance of epithelial architecture in LUAD, this may be overly reductive, neglecting other features such as the appearance of the stroma. We were motivated to map out the detailed spectrum of morphological heterogeneity in LUAD as a whole, ensuring we captured these underused morphological aspects. We used self-supervised learning (SSL) for this task, a form of AI that does not require up-front annotation, or any defined task, but in which the model is optimised to qualitatively encode meaningful histopathological features. SSL is particularly suited to this task, as learned embeddings can be used to identify latent phenotypes of potential clinical importance, enabling a data-driven re-exploration of existing schema. We used histomorphological phenotype learning (HPL) [12] to discover these recurrent morphological modules, building a disease-specific model, from a highly curated image set of over 4000 diagnostic LUAD H&E slides. This approach used real-world diagnostic images, and provided us with an accessible and interpretable suite of morphologies that we could subsequently link to clinical, pathological and molecular ground truths.

## Results

### Self-supervised learning gives us a topography of lung adenocarcinoma

The study design is summarised in Figure 1. The Leicester Archival Thoracic Tumour Investigatory Cohort - Adenocarcinoma (LATTICeA) dataset comprises 4427 tumour-containing slides from 1007 patients (Supplementary Table 1), amounting to over 8 m^2^ of glass slides and 15 TB of whole slide image (WSI) data. We trained an HPL model with these data, focusing on low magnification morphology (1.8 *µ*m per pixel; approximately equivalent to a 5x objective), motivated by the fact that tumour architecture is the most influential prognostic feature in LUAD [4]. We also made use of a linked physical resource of 23 tissue microarrays (TMAs), made from the same archival tissue blocks, containing up to three 1 mm cores per case giving approximately 3000 tissue specimens.

**Figure 1.**
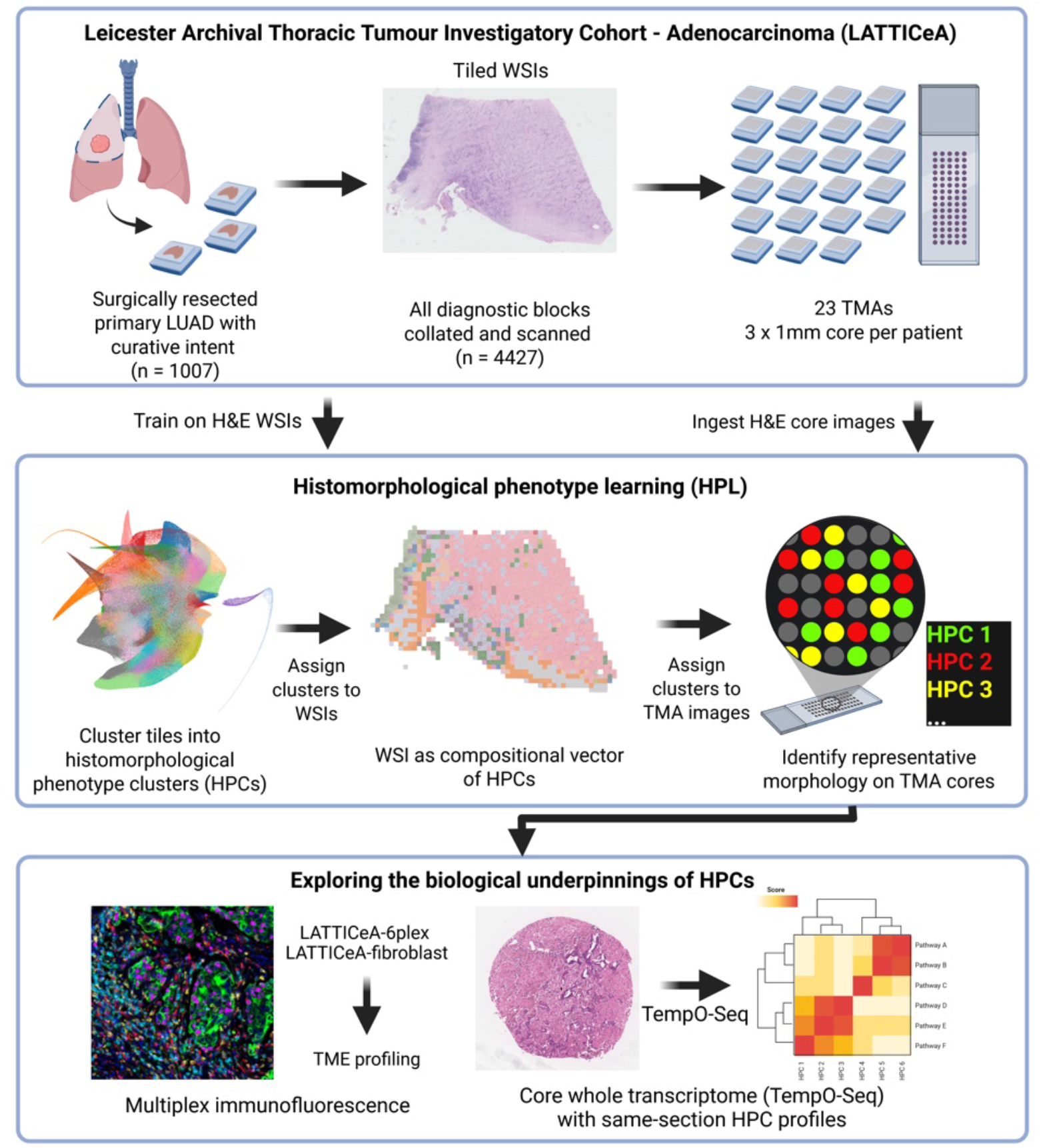
Overview of the study design. The Leicester Archival Thoracic Tumour Investigatory Cohort - Adenocarcinoma (LATTICeA) consists of over 1000 patients who underwent surgical resection of primary LUAD, with diagnostic WSIs, tissue on 23 TMAs and spatial proteomic and transcriptomic data. All diagnostic slides from each patient were scanned. WSIs are tessellated, and image tiles are used to train the self-supervised HPL model. Similar tiles are clustered, giving clusters of similar image tiles (HPCs). The frozen model was applied to H&E images of TMA cores, giving cores HPC identities which can then be associated with biological modules at the transcriptome and protein level. LUAD lung adenocarcinoma; WSI whole slide image; TMA tissue microarray; HPL histomorphological phenotype learning; HPC histomorphological phenotype cluster; H&E haematoxylin and eosin. Illustration created in BioRender.

We discovered 71 histomorphological phenotype clusters (HPCs), which we mapped to LATTICeA and the LUAD collection from The Cancer Genome Atlas (TCGA-LUAD). We also assigned HPCs to H&E images of TMA cores giving us physical examples of each HPC that we can use for downstream biological discovery. To give clinicopathological meaning to the AI-discovered HPCs, we obtained and integrated 3 pathologists’ annotations (Supplementary Tables 2, 3).

We wanted to ensure that no single HPC was dominated by contributions from a single patient. Each HPC can be considered as a vector of patient frequency (ie. each patient accounts for a certain proportion of the HPC). We examined the highest proportion of that patient frequency vector coming from a single patient on both the training set and TCGA-LUAD. We saw no HPCs in which the majority of the tiles came from a single patient, indicating satisfactory generalisability across patients and cohorts. We saw similar distributions in LATTICeA and TCGA-LUAD, reinforcing this point (Supplementary Figure S1a-d).

We first examined the image feature embedding space, looking for structure which aligns with existing histopathological ground truths (Figure 2a-d). We saw co-localisation of major tissue classes (Figure 2b). In addition to representation of well-recognised largely epithelially-defined growth patterns (Figure 2c, 2e), we found numerous HPCs defined by their stromal morphology, particularly in terms of the degree of inflammatory activity, as well as the stromal cellularity and fraction (Figure 2d, f).

**Figure 2.**
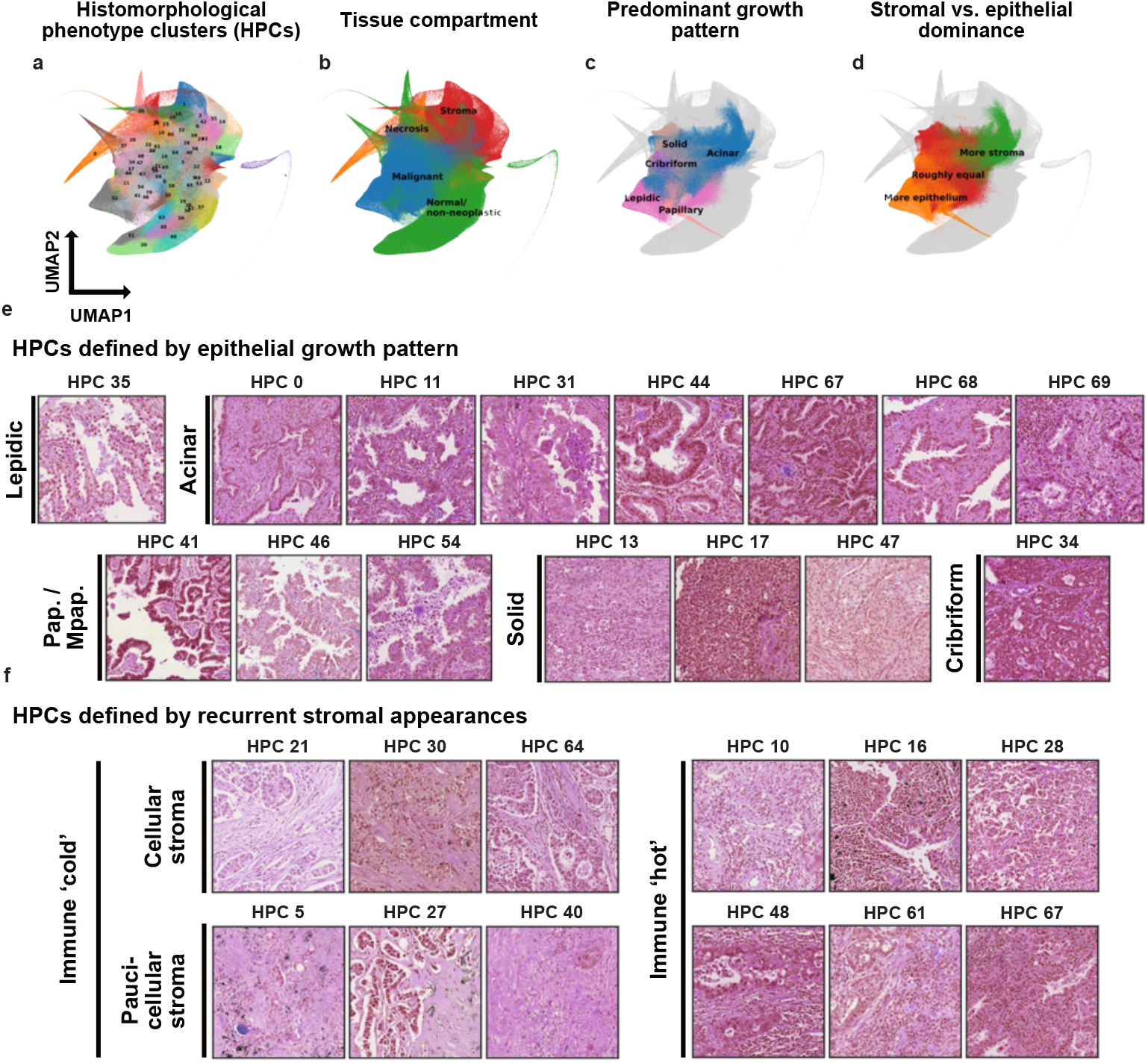
HPCs highlight variant stromal phenotypes and recapitulate classical growth patterns. (a-d) Uniform manifold approximation projection (UMAP) plots of the embedding space where each point is represented by a tile. Each tile is coloured according to its HPC identity. HPCs can be histologically reviewed and annotated according to their histological features, such as their parent HPC (a), the presence of malignancy (b), the predominant growth pattern (c) or stroma:epithelium ratio (d). (e) Example tile images showing that some HPCs are defined by consistent epithelial growth pattern. Pap. papillary; Mpap. micropapillary. (f) Example tile images of HPCs with variant stromal appearances, highlighting diversity in stromal morphology. All tiles H&E, widths 403.2 *µ*m.

We summarised all patients’ tumours as compositional vectors of HPC frequency, using the consensus-agreed malignant HPCs as the feature set, which allowed us to examine the relationship between HPC frequency and patient-level ground truths. We saw that HPCs representing certain morphologies associated with classical driver mutations, such as low-grade patterns with EGFR mutation and invasive patterns with TP53 mutation (Supplementary Figures S2a-c).

We next looked at the ability of these HPCs to predict survival (Supplementary Figure S3a). Reassuringly, the HPCs associated with the best prognosis were of IASLC low-grade growth patterns, and they also displayed high tumour-infiltrating lymphocyte (TIL) load (Supplementary Figure S3a, g). Interestingly, HPCs with the very worst prognostic implications combine being mostly stromal with being immunologically ‘cold’ (Supplementary Figure S3a, g), rather than by classical high-risk epithelial morphology such as solid pattern, although such clusters are indeed also related to poor outcome. The highest-risk HPC in our five-fold cross-validated survival model was HPC 27, which is predominantly acinar in growth pattern. Shapley Additive Explanations (SHAP) values corroborate this (Supplementary Figure S3b-d), which gave us an orthogonal measure of HPC contribution to the risk score at both the global and individual levels. Using risk scores derived from HPC abundance we could effectively stratify patients into risk groups (Supplementary Figure S3e, f).

Thus while current human grading depends upon scrutiny of epithelial morphology, we find that the addition of stromal appearances is, if anything, more prognostically powerful. To explore this idea further, we repeated survival modelling using the subset of HPCs that our pathologists identified as being defined by stromal morphology. These HPCs show tumour-associated stromal appearances such as collagenosis or elastosis but contain few, if any, epithelial elements or normal tissue. These diagnostically relatively neglected tumour areas accounted for 18 of the 71 HPCs, and 23% of modelled tissue area across the cohort. Of note, we found HPC 3 to be consistently associated with a poor prognosis (Supplementary Figure S4a-d, f), a cluster consisting of plump fibroblasts with interspersed collagen fibres. Conversely, areas of elastotic stroma with scattered lymphocytic aggregates (HPCs 56 and 60) were associated with more favourable prognosis.

We compared the relative prognostic power of our survival models built on malignant HPCs (‘HPL tumour’), stromal HPCs (‘HPL stroma’) and in combination (Supplementary Figure S4e). We also compared this to IASLC grade. We found that the most prognostic power came from the combined (tumour and stroma) model (train C-index mean *±* 1 standard deviation (SD) 0.655 *±* 0.018; test 0.638 *±* 0.04; and TCGA-LUAD 0.619 *±* 0.09), outperforming IASLC grade consistently (train C-index mean *±* 1 SD 0.585 *±* 0.012, test 0.586 *±* 0.044 and TCGA-LUAD 0.555 *±* 0.0). Interestingly, the performance of the stroma model was more stable across sets in each fold. This may indicate greater generalisability of stromal features, perhaps due to them representing recurrent physiological patterns of host tissue response rather than being the product of unique combinations of driver mutations, which underlies malignant epithelial appearances.

### Epithelial discohesion and T-cell infiltration as morphological axes

We noticed that several of the most lethal HPCs (such as HPCs 27, 40, 64 and 30) show marked infiltrative morphology superimposed upon classical growth patterns. This motivated us to explore the implications of epithelial discohesion so from this point on, we elected to focus on HPCs which were conventionally invasive, excluding those with lepidic, papillary and non-stromally invasive micropapillary growth patterns. To investigate this, we first identified cellular lineages in our TMA cohort by multiplex immunofluorescence (mIF) using a 6-plex phenotyping panel (Supplementary Figure S5) which gives us measures of cell density.

We quantified the degree of epithelial discohesion per HPC, using density-based spatial clustering of applications with noise (DBSCAN) (Figure 3a, see methods for further details). We hypothesised that the implications of epithelial discohesion depended on the stromal context, so we additionally measured T-cell density which allowed us to position each HPC along these two axes (Figures 3b-d). HPCs were immediately split by the immunologically ‘hot’/’cold’ axis by unsupervised hierarchical clustering alongside discohesion, which led us to group HPCs into superclusters based on these features, with each HPC being either ‘hot’/’cold’ and ‘cohesive’/’discohesive’. We validated these observations by calculating estimates of stromal cell counts from TCGA bulk RNASeq data, identifying correlations between the proportion of supercluster and cell counts (Figures 3e). Interestingly, there was a reduction in B cells in the high risk hot discohesive group compared to the other hot phenotypes, indicating a role for these cells to maintain a favourable immune response.

**Figure 3.**
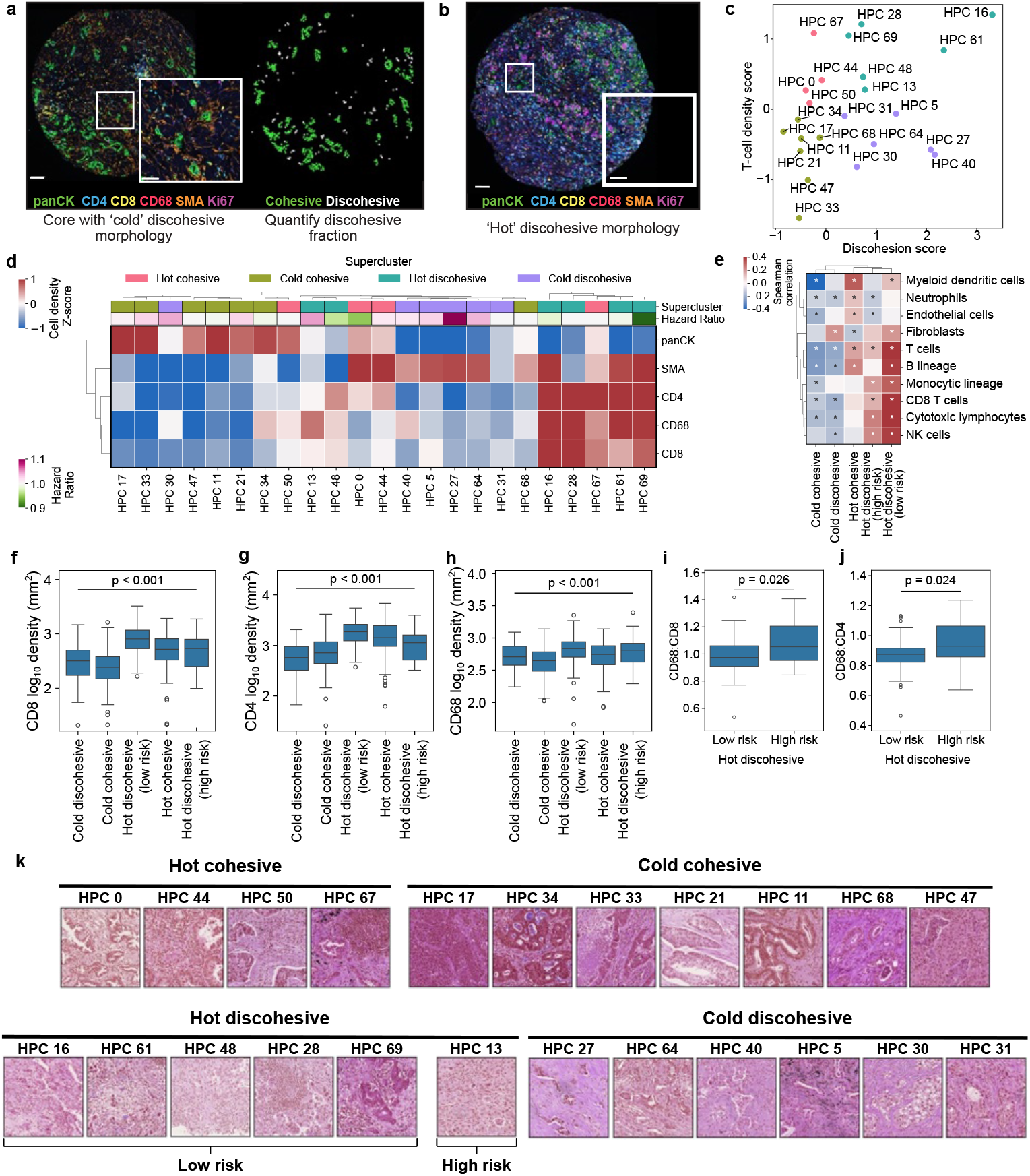
Epithelial and stromal morphology unite HPCs into superclusters with distinct cellular composition. (a) Graphical illustration of a core with discohesive morphology, and how DBSCAN isolates discohesive cells (white) from cohesive epithelial cells (green). Scale bars: main image 100 *µ*m, insert 50 *µ*m. (b) Example core with discohesive epithelial morphology and a dense T-cell infiltrate. Scale bars: main image 100 *µ*m, insert 25 *µ*m. (c) Discohesion score and T-cell density of each HPC, expressed as a Z-score. Each point is coloured according to its supercluster membership, as in (d). (d) Mean cell densities by HPC. Values are Z-scores, scaled by row. Each HPC is labelled with its average hazard ratio from the survival model and supercluster identity. (e) Spearman correlation between MCP Counter cell estimates and supercluster frequency from TCGA-LUAD (^*^ *p<* 0.05 after correction for multiple testing with Benjamini-Hochberg method), n=431. (f-g) *Log*_10_ density of CD8 T-cells (f), CD4 T-cells (g) and CD68 macrophages (h) by supercluster. Cold discohesive *n* = 105, cold cohesive *n* = 146, hot discohesive (low risk) *n* = 85, hot cohesive *n* = 127, hot discohesive (high risk) *n* = 19. (i-j) CD68 macrophage to CD8 T-cell (i) CD4 T-cell (j) *log*_10_ density ratios by hot discohesive supercluster. (k) Example H&E-stained image tiles of HPCs belonging to each supercluster. All tile widths 403.2*µ*m.

Cold cohesive HPCs were predominantly comprised of tumour cells, the cold discohesive group had an enrichment in SMA+ cells and the hot groups showed diversity in inflammatory cell makeup (Figure 3d). The cold discohesive HPCs were generally associated with a poor prognosis, and the hot discohesive with a more favourable one. Of the hot discohesive HPCs, HPC 13 has a negative association with prognosis despite belonging to the hot discohesive group. This HPC had a lower T-cell density than the rest of the hot discohesive HPCs but still more than either of the cold groups, suggesting that this difference in prognosis is not explained simply by a reduction in cytotoxic inflammatory activity (Figure 3f, g). The density of macrophages was comparable between the hot discohesive groups (Figure 3h), as such the ratio of macrophages to T-cells was increased in the high risk group (Figure 3i, j) suggesting a distinct microenvironmental recipe that drives this lethal phenotype. An example tile from each supercluster is shown Figure 3k,

### Epithelial discohesion and stromal inflammation predict survival

We next sought to identify the clinicopathological significance of these morphological axes. We went back to the WSIs, and expressed each whole tumour as a composition of superclusters. We then linked enrichment for superclusters with nodal status, pleural invasion and PD-L1 positivity. Tumours with a high fraction of cold discohesive HPCs were associated with node-positive disease at time of surgery (L2FC 0.320, *p<* 0.01) and pleural invasion (L2FC 0.287, *p<* 0.01) (Figure 4a, b) whereas hot cohesive disease shows the reverse. Both hot discohesive patterns were associated with PD-L1 positivity (low-risk L2FC 0.583, high risk L2FC 0.920, both *p<* 0.01; Figure 4c, Supplementary Figure S7a, b), though the relationship was stronger with the high risk group further distinguishing it as a distinct entity.

**Figure 4.**
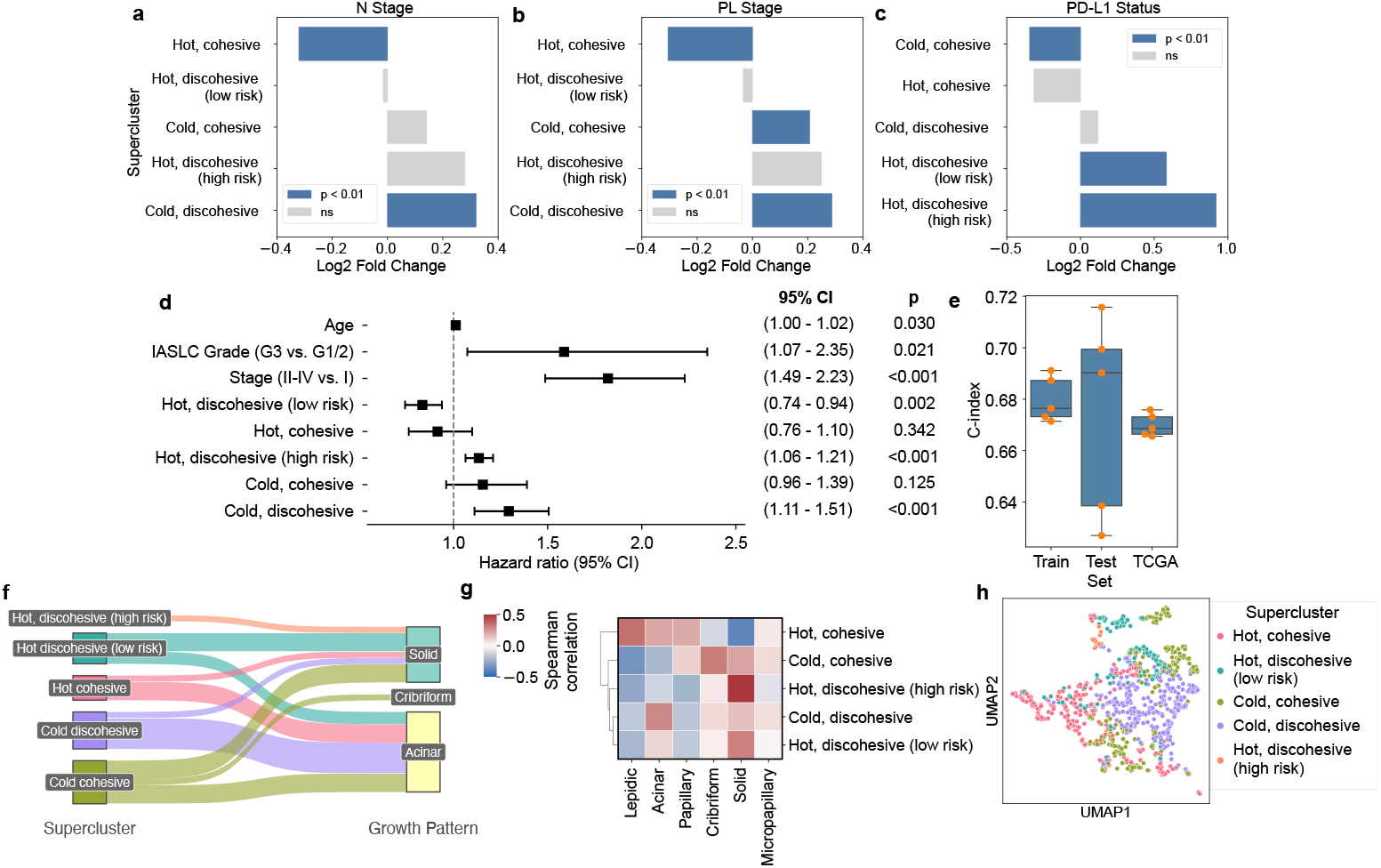
Cold discohesion identifies high risk acinar pattern. (a-c) Enrichment of supercluster frequency by nodal status; node-negative n=519, node-positive n=319, (a), by pleural invasion status; no pleural invasion n=446, pleural invasion n=456 (b) and by PD-L1 status; PD-L1 negative n=564; PD-L1 positive n=177 (c). Bars are coloured blue where where adjusted *p<* 0.01. (d) Multivariate Cox model using clinical and pathological parameters. (e) C-index from five-fold cross-validated multivariate Cox models. Box = IQR with median line; whiskers to *±* 1.5 × *IQR*. (f) Sankey plot showing the association between superclusters and growth pattern. (g) Correlation matrix showing the relationship between growth pattern and supercluster frequency at the whole tumour level. (h) UMAP plot of patients, coloured by the predominant supercluster.

In a cross-validated survival model, cold discohesive patterns were the most lethal of the five groups (hazard ratio 1.29, 95% CI 1.11-1.51, *p<* 0.001). High-risk hot discohesive morphology was also associated with lethality (hazard ratio 1.13, 95% CI 1.05-1.21, *p<* 0.001; Figure 4d). These remained prognostic alongside age, stage and grade in multivariate modelling. These models generalised to the unseen TCGA-LUAD cohort (Figure 4e).

To position superclusters alongside growth pattern, we returned to our histopathological annotations of each HPC which included the predominant growth pattern. These annotations were generated by examination of representative sets of small image tiles isolated from the remainder of the tumour, so to provide an orthogonal measure of the relationship between superclusters and growth pattern we also correlated the proportion of supercluster in the whole tumour against histopathologist whole-tumour growth pattern annotations. We found that almost all of the highly virulent HPCs which make up the cold discohesive group have predominantly acinar growth pattern and the low risk hot discohesive group predominantly solid (Figure 4f, g). This exemplifies the fact that morphology beyond grade can be scrutinised to identify high risk patients who may warrant more intensive treatment or closer follow up. Finally, by examining a UMAP plot, generated using each patient’s full HPC frequency vector, we see that patients with the same predominant supercluster occupy common niches in that plot (Figure 4h), suggesting distinct morphological ‘recipes’ for these five classes of tumour.

### Spatial proteomics defines twelve distinct cellular neighbourhoods

We further characterised the stromal composition of the superclusters to better understand their clinicopathological implications. We used our microenvironnment-profiling PhenoCycler panel (LATTICeA fibroblast, Supplementary Figure S8), focusing on a subset of stromal markers which highlight key cell types and states, including immune cell lineages, fibroblast phenotypes and extracellular matrix (ECM) proteins. We found the highly lethal cold discohesive regions to be enriched in COL1A protein expression (COL1A enrichment score 0.758, *p <* 0.01; Supplementary Figure S9a-b).

We next defined fibroblast phenotypes by unsupervised clustering of non-epithelial, non-immune cells. This generated six classes of fibroblasts (Figure 5a), characterised by their protein expression profile, and a single class of CD31-high endothelial cells. We used these detailed cell-level annotations for classical neighbourhood analysis, similar to that done in [13]. Briefly, this involves, for each cell, counting the number of cell types in a given radius then using unsupervised clustering to generate recurrent cellular neighbourhoods (CNs).

**Figure 5.**
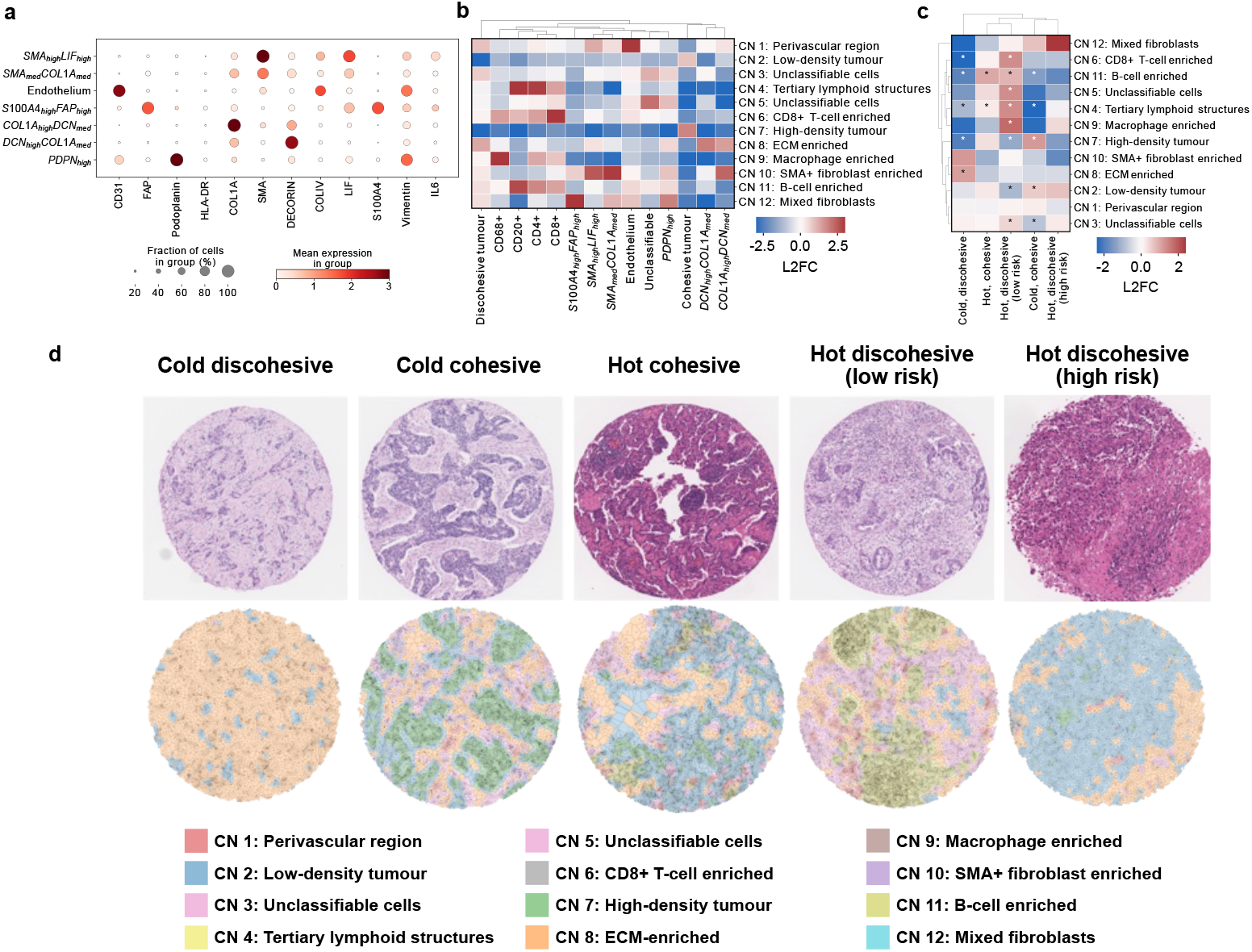
H&E superclusters are defined by cellular neighbourhoods. (A) Non-immune stromal phenotypes discovered with unsupervised clustering. The size of the dot is the fraction of cells of that phenotype which express the indicated marker, and the colour is the mean expression (fluorescence intensity) of the marker within cells of that phenotype. (B) Enrichment of cell populations in cellular neighbourhoods. defined as log2 fold-change (L2FC) increase in cell frequency over the population mean. (C) Enrichment of cellular neighbourhoods by supercluster; cold discohesive, n=44, hot cohesive, n=44, hot discohesive (low risk), n=50, cold cohesive, n=53; hot discohesive (high risk), n=8. (^*^ *p<* 0.05). (D) Example H&E stained images of superclusters and paired neighbourhood maps, where the region around each cell is coloured according to its neighbourhood membership.

We identified 12 distinct CNs, 8 of which represent distinct modes of stromal composition and immune cell infiltration (Figure 5).

### H&E superclusters are defined by cellular neighbourhoods

CN 8 was comprised of discohesive tumour cells, and both COL1A- and decorin-high matrix cancer-associated fibroblasts (mCAFs) representing an immunologically-excluded matrix-rich niche. Other fibroblast-enriched CNs show preference for SMA+ myofibroblasts and FAP+ fibroblasts (CNs 10 and 12 respectively). There were also inflammatory cell-rich CNs representing different classical modes of immune-cell activity (CNs 4, 6, 9 and 11; Figure 5b).

We then examined the frequency of neighbourhoods in TMA core images belonging to each supercluster (Figure 5c). As expected, cold discohesive cores were enriched for the matrix-rich CN 8 suggesting that there may be epithelial-stromal cross-talk facilitating this particular morphology and its clinical behaviour.

The high risk hot discohesive tumours areas are fewer in number, so there were no statistically significant relationships with CNs, but there was a trend towards FAP+ and SMA+ fibroblasts being more abundant in this group. Cold cohesive tumours, being related to solid pattern adenocarcinoma, were comprised mainly of tumour cells.

To more closely examine the stromal niche of the discohesive tumour cell, we carried out a further neighbourhood analysis focusing on superclusters with discohesive morphology. We calculated the stromal cell density as a function of distance from discohesive tumours cells in the three discohesive superclusters (Supplementary Figure S10a-c). At all distances, discohesive tumour cells in the cold discohesive supercluster tended to co-localise with mCAFs (Supplementary Figure S10a). In both hot discohesive superclusters, macrophages were the most abundant cell type near the discohesive population (Supplementary Figure S10b-c), but in the low-risk hot discohesive group these were accompanied by CD8+ T-cells and CD20+ B-cells, although not statistically significantly moreso in the high risk tumours over the low risk (Supplementary Figure S10d-h).

### Regionally resolved transcriptomics links cold discohesion to stromal remodelling

We examined the transcriptomic profiles which underpin these influential morphologies. For one TMA core section per case, we acquired whole-transcriptome data (TempO-Seq) from the same material which had been imaged for the HPL model, allowing for precise morphomolecular alignment. Inspection of a UMAP plot of the whole transcriptome showed broad associations between cores of common supercluster identity (Figure 6a).

**Figure 6.**
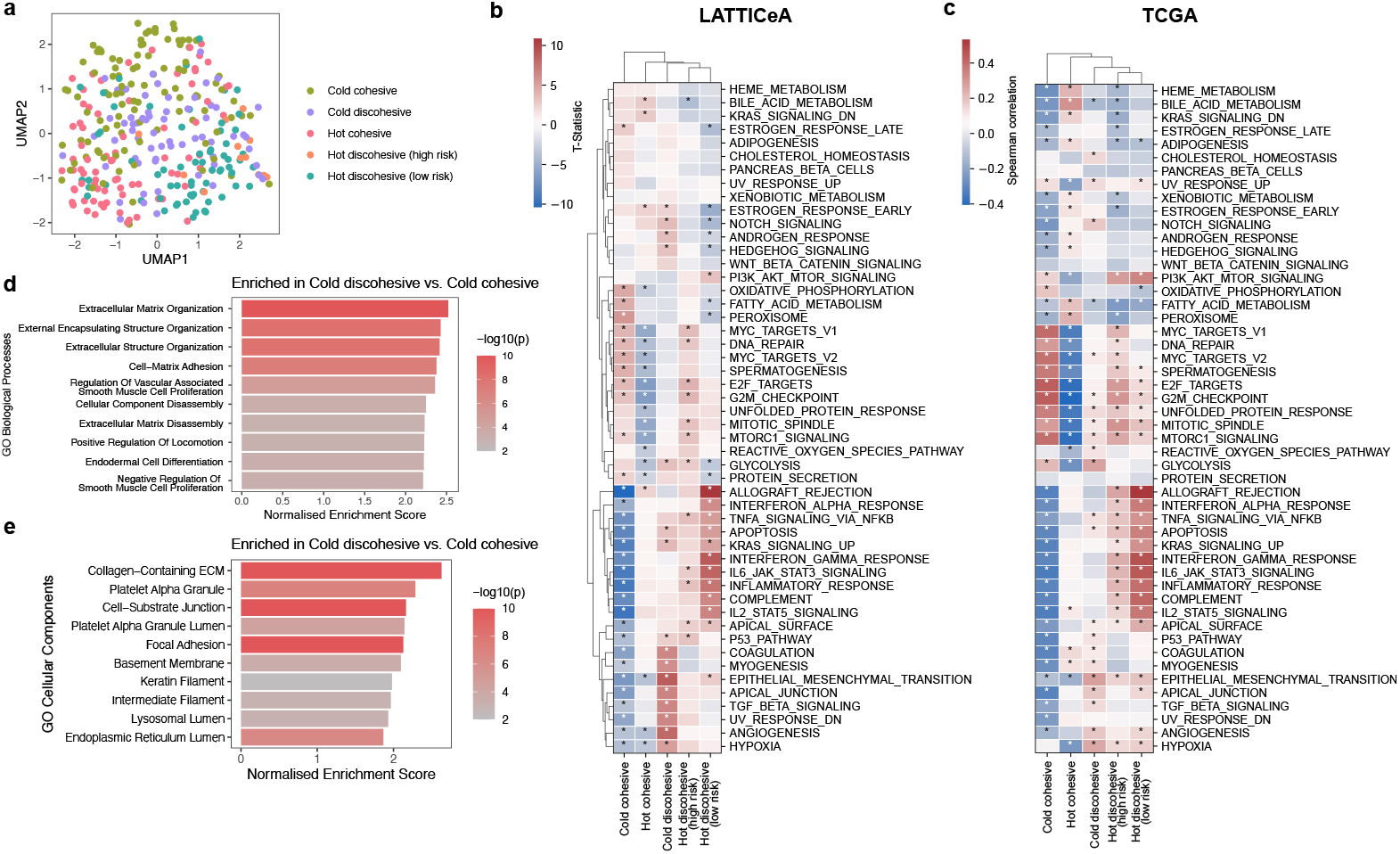
Cold discohesion is linked to stromal remodelling. (a) UMAP plot of LATTICeA cores coloured by core supercluster. Cold cohesive, n=100; cold discohesive, n=63; hot cohesive, n=76; hot discohesive (low risk), n=65; hot discohesive (high risk), n=13. (b) Enrichment of ssGSEA scores (MSigDB hallmarks database) on LATTICeA. (c) Correlation of supercluster fraction against ssGSEA enrichment score on TCGA-LUAD, n=431; ^*^ *p<* 0.05. (d) Geneset enrichment analysis on differentially expressed genes between cold discohesive and cold cohesive cores on LATTICeA (GO cellular components). (e) Gene-set enrichment analysis on differentially expressed genes between cold discohesive and cold cohesive cores on LATTICeA (GO biological processes)

Single-sample gene set enrichment analysis (ssGSEA) showed that cold discohesive tumour regions have a gene expression programme directing invasion, via epithelial-mesenchymal transition (EMT) and stromal remodelling (angiogenesis, myogenesis, TGF-beta signalling), and hypoxia (Figure 6b). This is suggestive of a highly motile autonomous population which actively remodels its local environment, possibly to facilitate metastasis. We made similar observations by using TCGA-LUAD bulk RNASeq data (Figure 6c), confirming relationships between these pathways and the proportion of cold discohesive morphology.

To further highlight the virulent programmes of discohesion, we compared gene expression profiles between cold discohesive and cold cohesive morphologies. On gene set enrichment analysis (GSEA), showed that the cold discohesive morphology is related to pathways such as collagen-containing extracellular matrix and extracellular matrix organisation, agreeing with our protein-level findings (Figure 6e, f). This is accompanied by upregulation of cellular adhesion to the ECM, and with ECM disassembly programmes, indicative of active stromal remodelling. We speculate that these findings reflect autonomous ECM-directed migration, explaining their metastatic potential.

## Discussion

We used SSL to discover, *ab initio*, the defining set of H&E morphologies that comprise LUAD, using a vast image dataset of over 4000 WSIs from over 1000 patients. We identified highly prognostic morphological signatures, revealing the fact that stromal morphology is just as influential in predicting patient outcome as epithelial appearance. We go on to augment these H&E entities with detailed multi-modal molecular characterisation, at both the protein and RNA levels.

We identified epithelial cohesiveness and T-cell infiltration as crucial determinants of prognosis. It is notable that neither of these feature in current prognostic schema. Both are predictive of survival, additively to stage and grade, with a strikingly high risk profile seen in cold discohesive tumours. Tumour budding is the specific phenomenon of discohesive epithelial elements at the invasive front of a tumour [14], and has been associated with a poor prognosis in colorectal carcinoma (CRC) [14], LUAD [15], as well as other solid malignancies [16–18]. Although our definition of a discohesive island is modelled after the definition of a tumour bud, we describe discohesive patterns of growth throughout the tumour bed. The invasive front is not as clearly defined in LUAD as in other carcinomas, such as CRC, as these tumours often have a remodelled central stromal core which is often a focus of invasive activity [19, 20]. As such, we consider this morphology to be more akin to a growth pattern rather than the localised finding budding is currently commonly described as. A discohesive pattern of growth has been described in LUAD [21], and was found to be associated with poor prognosis. Similarly, in a study exploring automated grading in LUAD, Pan et al encountered a high risk variant of acinar pattern LUAD, acinar scattering, a metric of spatial dispersion of acinar islands throughout the tissue section [22]. However, these studies stop short of highlighting stromal context as the key determinant of the lethality associated with discohesive growth. Our work goes on to demonstrate that the lethality associated with epithelial discohesion is contingent on the paucity of T-cells, and their abundance redefines discohesion as a marker of more favourable prognosis.

Stromal morphology is influential in prognosis in multiple solid malignancies, such as the tumour-stroma ratio in colorectal cancer [23] and the degree of lymphocytic infiltration in melanoma [24]. Cancer-associated fibroblasts (CAFs) are heterogenous and lack a consensus molecular classification system, although several have been proposed [25–27]. We found the lethal cold discohesive morphology to be enriched for mCAFs, similar to those described by Cords et al [28]. In that study, patients with LUAD were grouped according to their fibroblast profile and those with an enrichment of mCAFs were associated with the worst prognosis. We associate these mCAFs with a consistent and readily identifiable phenotype rooted in H&E images which allows us to contextualise this information in real world histopathology. This critical step paves a way forward to enabling the integration of stromal profiling into conventional morphological assessment of LUAD for clinical decision making.

COL1A is the most abundant collagen, and its excess deposition is associated with poor prognosis in LUAD [29]. Once again, we make this observation anchored in features that are readily discernable in standard diagnostic histopathology, rather than dependent on a further molecular assay. CAFs, as the major producers of ECM proteins, exhibit mechanical pressure on invading tumour cells which can be anti-tumourigenic in nature [30]. However, the arrangement of collagen fibres in the ECM can modulate the invasive ability of the malignant population, with parallel arrangement of fibres facilitating cellular migration [31, 32]. Furthermore, this mechanical pressure on the invasive population can induce persistent transcriptional reprogramming resulting in a pro-migratory and pro-survival phenotype via activation of myosin II and STAT3 activity, leaving them primed for metastatic potential [32]. We observed transcriptomic modules for ECM remodelling and cellular migration enriched in cold discohesive tumours, possibly reflecting similar phenomena. It is unclear whether the excessive ECM deposition is initially caused by an aberrant host physiological response, which selects for a migratory population capable of traversing it, or if the invasive epithelial population induces such a response of its own accord. This would be a fascinating avenue for functional study as it raises the possibility of a CAF-directed therapeutic approach to mitigate metastatic dissemination in the neoadjuvant setting.

Our study has limitations. Firstly, our cohort is composed of patients who underwent surgery with curative intent so we lack representation of advanced clinical stage. Secondly, we only use resection material so we lack validation in diagnostic small biopsy material. Thirdly, in our immune microenvironment characterisation we focus on basic inflammatory cell phenotypes so we have not yet incorporated the role of the myeloid lineage or of macrophage polarisation. Finally, although our transcriptomic data are perfectly morphologically aligned and microregional (ie. from a 1 mm core and imaged in H&E prior to sequencing), they are still ‘bulk’ data, as is the validation dataset from TCGA-LUAD.

In conclusion, we did an unsupervised exploration of the morphological landscape of LUAD from an archival cohort of over 1000 patients, validating our observations in an independent set. We demonstrate that AI is not simply a tool for automation or efficiency gains in histopathology but that it can be a powerful research tool aiding biological discovery and offering new knowledge which can be immediately applied to refine clinical diagnostic definitions. We have constructed a foundational morphological dictionary of LUAD, applicable to any study involving diagnostic human LUAD tissue. This model is freely available to use for inference, which democratises the interpretation of the complex morphology in this disease, and offers a way to link rich multi-omic data back to real world diagnostic histopathology.

### Image datasets

The training set - the Leicester Archival Thoracic Tumour Investigatory Cohort, Adenocarcinoma (LATTICeA) - comprised 4427 WSIs from 1007 LUAD patients (Supplementary Table 1), with associated clinical annotation, who underwent surgical resection with curative intent from a single centre [20]. WSIs of H&E stained sections of 23 TMAs were acquired identically to the resection specimens.

The external test set was the LUAD collection from TCGA (TCGA-LUAD), which was retrieved as described in [12], which resulted in 501 WSIs from 439 patients. Clinical annotations were downloaded from GDC Portal. IASLC grade was retrieved from written pathology reports where possible, and elsewhere was assigned by reviewing the pathology slides directly (KR and JCL).

### Self-supervised learning and clustering

We used HPL which has previously been shown to identify recurrent morphological features in large pan-cancer image sets from TCGA [12]. The model uses Barlow Twins [33] to train a convolutional neural network with a ResNet backbone. Each tile is duplicated, with the pair being subject to augmentations, such as cropping, rotating, or modifying the colour. The aim is therefore to produce a representation from each tile in the pair where each element of the representation perfectly correlates with itself.

The self-supervised workflow was completed as follows:

- WSI pre-processing: We tessellated all images at 1.8 *µ*m per pixel, approximately equivalent to 5x magnification. WSIs of H&E-stained TMA sections were dearrayed using QuPath v0.5.1 [34] at native resolution (0.2523 *µ*m per pixel). Tiles were captured from core images, then downsampled to 1.8 *µ*m per pixel. Given the large size of the desired tiles (diameter approximately 400 *µ*m), we could only obtain 4 tiles from a single 1 mm core of tissue so we centred these on the centroid of the image.
- Self-supervised encoder: We trained the self-supervised model for 10 epochs with a batch size of 64. All hyperparameters were as previously described in Quiros et al [12]. The weights were frozen, and representations were obtained for all tiles in LATTICeA, TCGA-LUAD and the TMA set.
- Clustering: We subsequently used Leiden community detection to discover HPCs, which are groups of tiles with similar morphology. We initially do this at a high Leiden resolution (5.0) to identify and remove tiles with artefact such as areas where the image is blurred or out of focus. We removed these tiles and subsequently reperformed clustering at a lower Leiden resolution (2.5). We used the Leiden clustering implementations in scanpy v1.10.1 and rapids single cell v0.10.1 [35, 36]. We discovered clusters using 2.5 million randomly sampled tiles and assigned these clusters to all remaining tiles across all datasets.
- Mapping HPCs to the TMA set: As there were only 4 tiles per tissue core, we opted to use majority vote to classify whole cores as belonging to an HPC. We considered cores with ≥ 3 identical HPCs as ‘pure’ cores and included these in our analysis. Where the analysis is supercluster-based, if ≥ 3 HPCs all belonged to the same supercluster then the core was assigned to that supercluster even if there was no HPC majority (see Supplementary Figure S6a).

### Cluster interpretation

For each HPC, we randomly extracted 100 tiles from that HPC for review. We issued the same tile set to 3 pathologists (JLQ, DD, and KR) and collected quantitative and qualitative metrics via a standardised questionnaire. We collated numeric measurements by mean and categorical measurements by majority vote (Supplementary Tables 2, 3).

### Clinicopathological statistical analysis

For subsequent statistical testing, HPC frequency was counted as a proportion of all tiles present (ie. expressed as proportions). Where necessary, these numbers were projected out of Atchisonian space using a centered log ratio transformation prior to model construction using the clr function in scikit-bio 0.5.9 [37].

We carried the survival analysis in two steps. Firstly, we used cluster frequencies as our feature matrix, using the consensus-agreed malignant, stromal and all HPC sets. We trained Cox proportional hazards models in lifelines v0.26.3 [38] using overall survival as the event. Given the high number of features, we fit L1-regularised models. SHAP values were calculated using shap v0.44.1 [39]. For risk dichotomisation, each patient’s partial hazard was calculated and this was cut at the median to create high and low risk groups. This was done independently for the test set and TCGA-LUAD. We performed 5-fold cross-validation for the survival models, using TCGA-LUAD as an external test set for each training and validation split. For the supercluster survival models, HPC frequencies were aggregated by supercluster, and these features were used in the survival model. Baseline clinical variables were also included, namely age, stage (as a binary variable, stage I and stages II-IV) and IASLC grade (also as a binary variable, grades 1/2 and grade 3).

Categorical clinical labels (N-stage, PL-stage and PD-L1 status) were treated as binary labels for the enrichment task. The log2 fold change was calculated between the patients in the positive group against the overall population mean.

Unless stated otherwise, statistical significance between two groups was tested with the Mann-Whitney-U test and between multiple groups using a one-way ANOVA. Where appropriate, correction for multiple testing was done using the Benjamini-Hochberg method. All statistical tests were implemented in scipy v1.14.0 [40].

### Multiplex immunofluorescence

#### LATTICeA 6-plex

The multiplex assay (consisting of antibodies against CD68, pan-cytokeratin (AE1/AE3), SMA, CD4, CD8 and Ki67; with 4’,6-diamidino-2-phenylindole [DAPI] as a nuclear counter-stain) was applied to 23 4 *µ*m thick FFPE sections from the LATTICeA cohort using the Ventana Discovery Ultra autostainer (Roche Tissue Diagnostics, RUO Discovery Universal v21.00.0019). Discovery Cell Conditioning 1 (CC1) (Roche Tissue Diagnostics, 950-123) was applied to the section for 64 minutes at 95 °C for antigen retrieval. The details of the antibody application protocol have been described previously [41].

Image analysis was carried out in Visiopharm. Tissue segmentation was performed using DeepLabv3+ (v2023.01.3.14018), after training on annotations done by a pathologist using DAPI, cytokeratin and autofluorescence as the input. The result was a probability of each pixel belonging to tumour, stroma, necrosis or background regions and these were assigned when the feature probability exceeded 0.5 (50%).

For cell detection, a separate deep learning algorithm based on the U-Net architecture (version 2022.12.0.12865) was trained using the DAPI input channel to identify background, boundary, and nuclear features. Nuclear, boundary, and background labels were assigned when feature probabilities surpassed the 0.5 threshold. Post-processing steps were applied to remove background and boundary labels. Cytoplasmic regions were generated by dilating nuclear labels—by 20 pixels for cells in tumour regions and 10 pixels for cells in stromal regions. Where cells overlapped, the boundary feature heatmap was used to separate cells. For each core image, output variables exported were: area, mean marker intensities, and X/Y coordinates of each cell.

#### LATTICeA fibroblast

All antibodies used in the panel (4) were validated in 3,3’-Diaminobenzidine (DAB) staining on the Ventana Discovery Ultra (Roche Tissue Diagnostics, RUO Discovery Universal V21.00.0019) using human FFPE control tissue known to express the marker of interest. After the optimal dilution was apparent, we conjugated each antibody (BSA free and azide free), guided by our modified in-house version of the Akoya PhenoCycler Conjugation protocol (5).

Four TMA sections were cut at 4 *µ*m and baked in a 60 °C oven for a minimum of 30 minutes. Each section was placed on the Roche Ventana Discovery ULTRA where antigen retrieval took place using CC1 (Roche Tissue Diagnostics, 950-123) at 95 °C for 32 minutes. Manual staining of antibodies occurred following User Guide [42]. Antibodies were applied as cocktails in 2 parts (6): The first application was applied for 3 hours at room temperature and the second round of antibody cocktail was applied overnight at 4 °C.

Akoya Reporters were later applied following the protocol template (Supplementary Figure S11), with an adjustment to the nuclear stain (10 *µ*l per well at 5 ms). Exposure times were kept at 150 ms for each marker and the experiment was run at 8-bit. Images were quality checked using QuPath v0.5.0.

Tissue segmentation was carried out by training a random trees pixel classifier in QuPath v0.5.1 using tumour, stroma and background classes. Areas of necrosis were manually excluded. Cell segmentation was done using Cellpose v3.0.1 [43] using the QuPath extension v0.9.6 [44] using the Cyto3 pre-trained model. Regional (tumour, stroma) marker fluorescence intensities were exported using the ‘add intensity features’ function using the default parameters. The mean cell fluorescence intensities were added using the Cellpose QuPath extension at time of cell segmentation.

#### Normalisation and phenotyping

To mitigate any batch effect, we scale all raw cell values per slide and per channel using Z-score.

For the 6-plex panel, we used unsupervised clustering using all markers to assign phenotypes to cells (tumour cell, CD4+ T-cell, CD8+ T-cell, CD68+ macrophage, SMA+ cell; Supplementary Figure S5). We projected cell labels back onto the original images with a pathologist (KR) examining 20% of the results and adjusting the phenotypes where required.

For the fibroblast panel, we also used unsupervised clustering to initially identify broad lineages of cells and collapse these into three main groups (tumour, immune, non-immune stromal). The subset of markers used in this study is specified in Supplementary Figure S8. Subsequently, we re-clustered to assign specific identities (eg. CD8+ T-cell) to these cells. Specifically for the fibroblast phenotyping task, we used K-Means clustering with k=9 (as implemented in scikit-learn v 1.4.0 [45]) using only the following channels: CD31, FAP, podoplanin, HLA-DR, COL1A1, SMA, decorin, COLIV, LIF, S100A4, vimentin and IL6. This was done on cells that were non-epithelial and non-immune in phenotype (ie. which remained without an identity after the previous round of clustering).

### Defining superclusters

To define epithelial discohesion, we took inspiration from the criteria used in the reporting of colorectal carcinoma [14]. We used density-based spatial clustering of applications and noise (DBSCAN), as implemented in scikitlearn v1.4.0, using eps=25 and min samples=5 on the tumour cells on the 6-plex panel. This clustering algorithm assigns points in 2D space to a cluster if they meet specified minimum criteria for number of cells (min samples) in a given radius (eps). We required a less than 5 cells in a 25 *µ*m radius to call a discohesive cluster, termed ‘noise’ points in the DBSCAN lexicon. We calculated the proportion of noise cells of all tumour cells to measure epithelial discohesion. We calculated the mean noise proportion of all cores and used that as the cut-off to define cohesion/discohesion. The discohesion score for each HPC was calculated as the mean of the discohesion score of all pure cores belonging to that HPC. The cut-off was then applied to categorise HPCs as cohesive/discohesive. A similar approach was used for T-cell density (‘hot’/’cold’). The density of T-cells was the sum of the densities of CD4+ and CD8+ T-cells and subsequently log-10 scaled. Again, we calculated the mean of all pure cores to establish a cut-off then calculated the mean T-cell density per HPC and used the cut-off to classify HPCs. The number of cores and their HPC identities used in this calculation is specified in Supplementary Figure S6b.

Given we asked questions of the interaction between invasive epithelial architecture and the composition of the tumour microenvironment, we did not consider HPCs 35, 41, 46 and 54 in the classification. HPC 35 is representative of lepidic growth pattern, a low-grade non-invasive pattern. Similarly, HPCs 41, 46 and 54 show papillary and micropapillary growth, characterised by upward projections of tumour into the airspace rather than stromal invasion.

### Neighbourhood analysis

We defined neighbourhoods based on the counts of cells within 25 *µ*m radius of the index cell. We calculated these using the KDTree function from scipy. We then processed these using K-Means clustering, with k=6 using scikitlearn. To characterise the neighbourhoods, we calculated the log2 fold change (L2FC) of cell densities of all cell types relative the population mean. Similarly, we calculated the mean frequency of each cellular neighbourhood per supercluster and expressed this as L2FC from the population mean. Neighbourhood frequency was expressed as a proportion of all neighbourhoods per core. Enrichment scores in region- and cell-based analyses were calculated by subtracting the mean of the group of interest from the population mean.

### RNA-Seq analysis

The preparation of the LATTICeA TempO-Seq sequencing was as described previously [46]. Differential expression was done in DESeq2 v1.40.2 [47] between the cold discohesive and cold cohesive groups after excluding genes with a mean expression of *<* 1. GSEA was done in fgsea v1.26.0 [48] using the GO biological processes and cellular components gene sets. Pathways with an adjusted p-value of *<* 0.01 were considered significant. ssGSEA was done in GSVA v1.48.3 [49] using the MSigDB Hallmarks gene sets. The enrichment for ssGSEA scores at supercluster level was done in a one-versus-the-rest fashion, with significance tested with Welch’s t-test.

TCGA-LUAD RNASeq data was obtained from Xena browser [50] in *log*_2_*TPM* +1 format. Cell counts were estimated using MCP-counter v1.2.0 [51]. Correlation analyses were done against supercluster frequency for that whole tumour (ie. an aggregation of all slides available for a single patient).

### Tumour genotyping

Tissue acquisition and sequencing has been described previously [41]. In brief, 1 mm cores of tumour tissue were sampled for DNA sequencing at the time of TMA construction. DNA extraction was done using the QIAmp FFPE DNA tissue kit (Qiagen) according to the manufacturer’s instructions and quality checked with an Agilent Tapestation 4200 with Genomic screen tape. Sequencing was done using an ISO 15189 accredited custom Ampliseq based assay (MGP-4-DNA, Sarah Cannon Molecular Diagnostics). Library preparation was performed manually with templating and chip loading on an Ion ChefTM and sequencing on an Ion GeneStudioTM S5 Prime. Data analysis was carried out with Torrent Suite Variant Caller (5.10.1.20) combined with in-house developed applications for error checking, quality control, variant ‘binning’ and HGVS nomenclature construction. The assay was validated with a limit of detection for SNVs and small indels of 2.5% variant allele frequency in hotspot regions from *>*30 genes. Samples which failed to achieve full panel coverage or attain other quality control criteria were excluded.

## Code and Data Availability

The original HPL codebase is found on GitHub [12]. Modifications to support this manuscript, including all analysis code and instructions to map HPCs to external data can be found at https://github.com/K-Rakovic/HPL-LATTICeA. Model weights, reference HPCs required for inference, and source data will be made available on publication.

Some LATTICeA study data and materials (diagnostic H&E images and clinical annotation) are currently subject to a material and data transfer agreement between the University of Leicester, the University of Cambridge and NHS Greater Glasgow and Clyde. Restricted access data can be accessed by application to NHS Greater Glasgow and Clyde Biorepository as custodians; the data access request will be reviewed and released under their research ethics committee-approved tissue bank protocols. Requests will be reviewed and approved within 6–8 weeks and will be accompanied by a data sharing agreement detailing the conditions and restrictions of use and publication.

## Acknowledgements

The authors would like to extend their gratitude to: the NHS Greater Glasgow and Clyde Biorepository for their ongoing management of the LATTICeA cohort; to the CRUK Scotland Institute Deep Phenotyping Advanced Technology Core Facility (Research Resource Identifier [RRID]: SCR 027366) for preparation of mIF images; Naveed Khan, CRUK Scotland Institute for the management of the high performance computing facility; Catherine Winchester, of the CRUK Scotland Institute, for critically appraising this manuscript; and and Philip Bennett from Sarah Cannon Molecular Diagnostics, Part of HCA Healthcare UK for the preparation of the LATTICeA genomic dataset.

## Author Contributions

KR designed and executed the experiments, and prepared the manuscript. AP prepared the transcriptomic data and contributed to its analysis. ACQ provided the initial codebase and supervised its modification and implementation. ZH analysed the genomic data. JCL, DAD and CD provided histopathological expertise. DAM developed the LAT-TICeA dataset and MS, CW, AT and CF collated the data, built the TMAs and scanned the diagnostic images. SM, RLB, LOJ and IP developed and applied the mIF assays, and carried out image analysis. DC designed the fibroblast mIF panel and provided input into experimental design and interpretation. CJM provided computational resources and input into experimental design. KY and JLQ supervised the work. All authors reviewed and contributed to the final manuscript.

