## Supplementary Material for "Self-supervised AI reveals a lethal discohesive phenotype in lung adenocarcinoma"

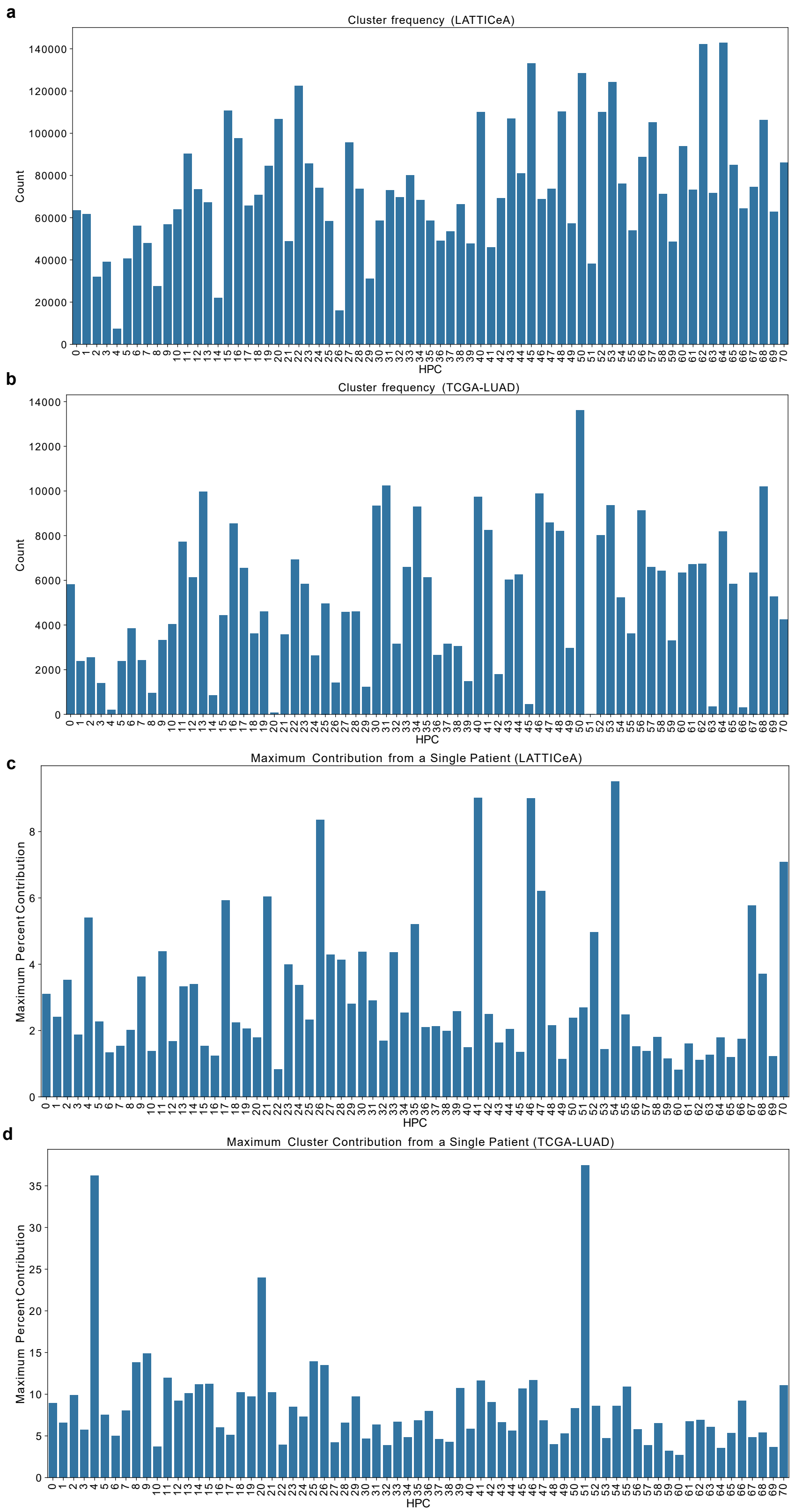

Supplementary Figure S1: (a, b) Cluster frequency in LATTICeA (a) and TCGA-LUAD (b). (c, d) The proportion of an HPC coming from a single patient.

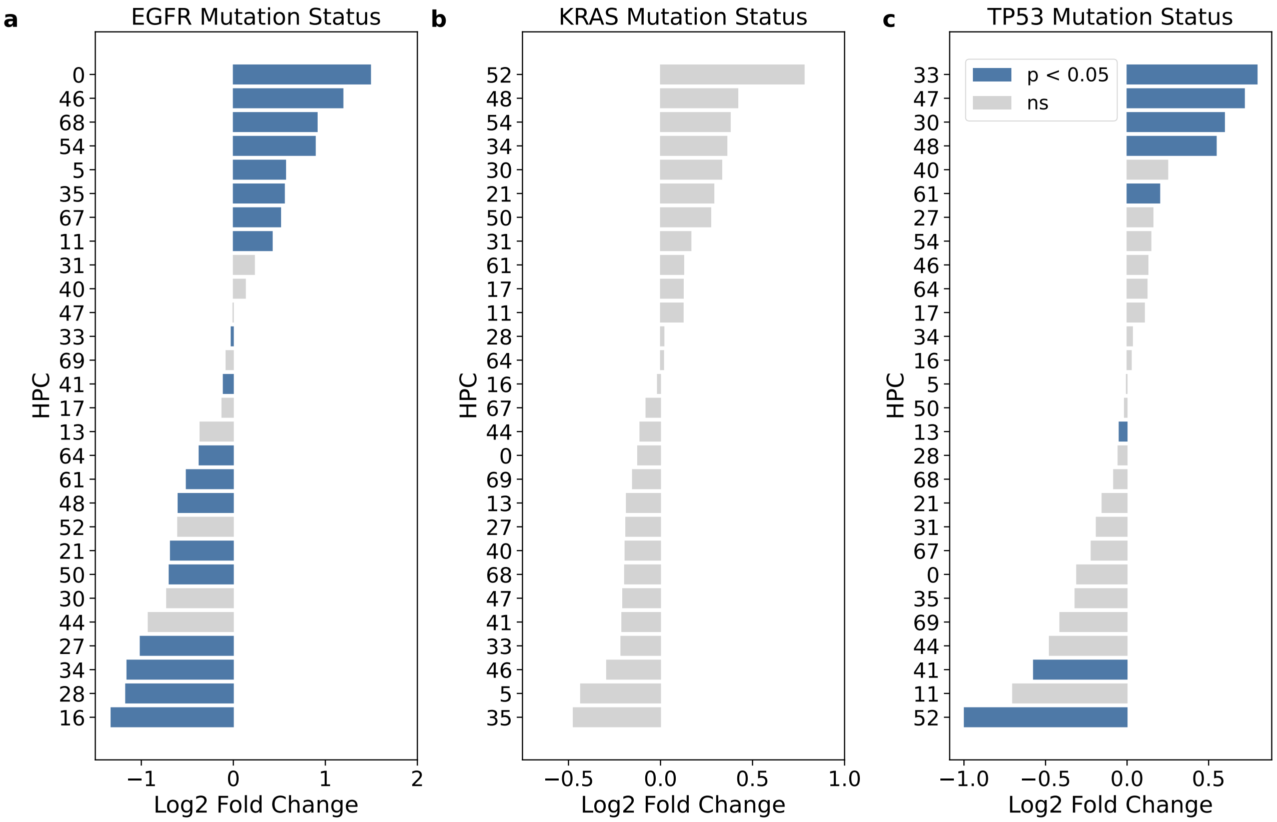

Supplementary Figure S2: Enrichment of HPCs in cases which are EGFR-mutant n=80 WT n=564 (a), KRAS-mutant n=256 WT n=388 (b) and TP53-mutant n=225 WT n=419 (c) compared to wild type. Statistical significance tested with a Mann-Whitney-U Test. Coloured bars show HPCs for which are statistical significance is maintained ($p < 0.05$) after correction for testing with Benjamini-Hochberg method.

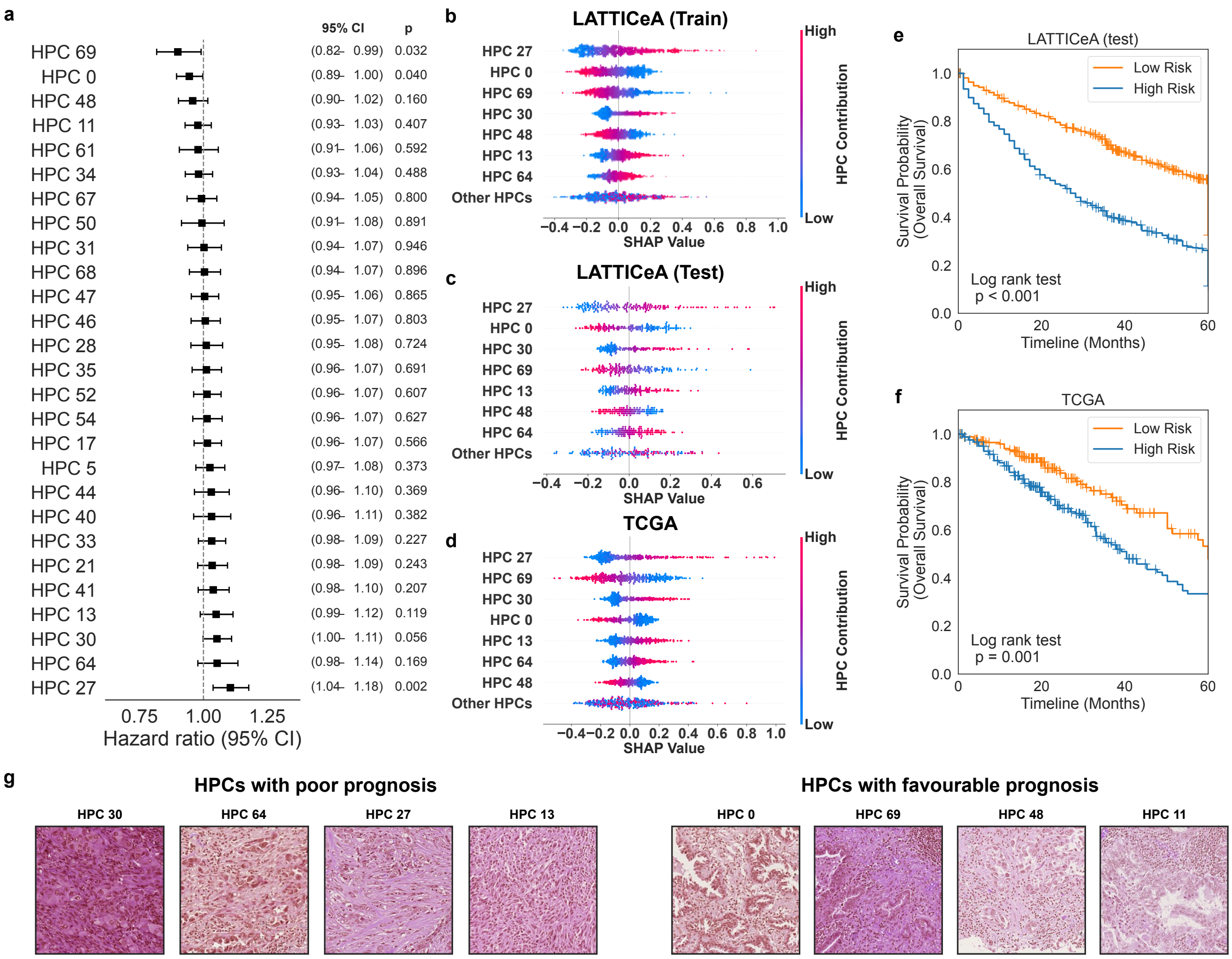

Supplementary Figure S3: (a) Forest plot summarising the Cox proportional hazards model trained on the HPC frequency vector of malignant tumour HPCs. (b-d) SHAP values for the train (b), test (c) and TCGA (d) sets derived from predictions calculated by the survival model. (e, f) Kaplan-Meier plots of the LATTICeA test set (e; low risk n=467, high risk n=474) and TCGA-LUAD (f; low risk n=174, high risk n=244), dichotomised according to their predicted survival. (g) Exemplar tile images of the four HPCs with the lowest and highest hazard ratios. HPCs associated with favourable outcome are predominantly low grade with frequent lymphocytes, whereas poor outcome HPCs are characterised by cold collagenous stroma. Tile widths 403.2 $\mu$m.

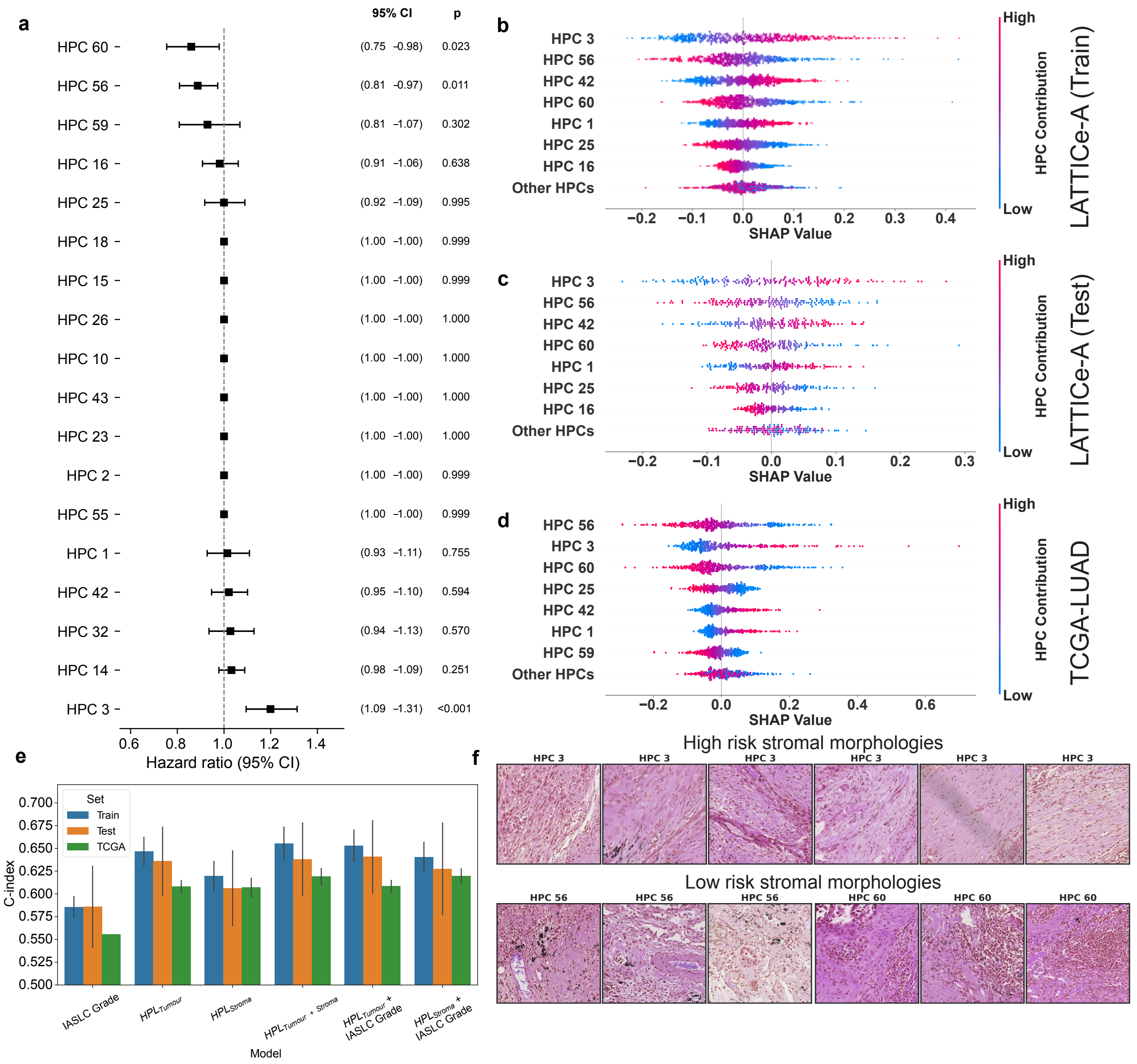

Supplementary Figure S4: (a) Forest plot summarising the Cox proportional hazards model trained on the HPC frequency vector of abnormal stromal HPCs. (b-d) SHAP values for the train (b), test (c) and TCGA (d) sets derived from predictions calculated by the survival model. (e) Mean c-index across five folds calculated for the train, test and TCGA sets using different sets of features: IASLC grade alone, tumour and stroma HPCs alone, in combination with each other and with IASLC grade. Error bars are $\pm$ 1 standard deviation. (f) Exemplar tile images of HPC 3, associated with poor prognosis, and HPCs 56 and 60, associated with more favourable prognosis. Tile widths 403.2 $\mu$m.

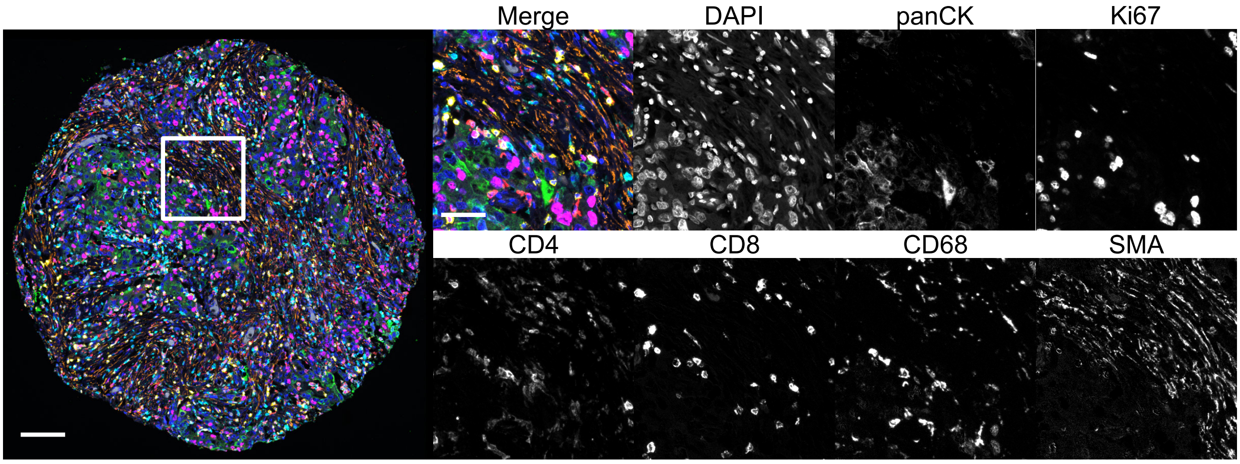

Supplementary Figure S5: Example mIF image from the 6-plex panel (see Figure 3). Scale bars: core 100 $\mu$m, inserts 30 $\mu$m.

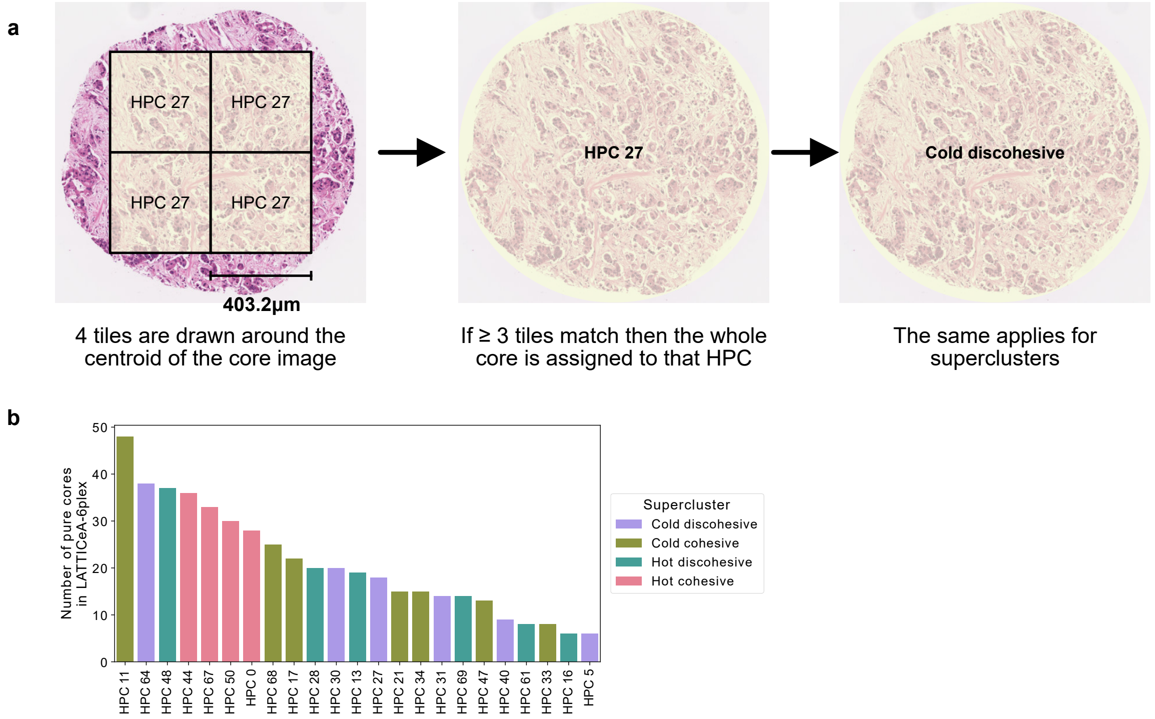

Supplementary Figure S6: (a) Schematic illustrating how HPCs are assigned first to tiles, and majority vote classifies the whole core are belonging to an HPC or supercluster. (b) Histogram of pure cores by HPC.

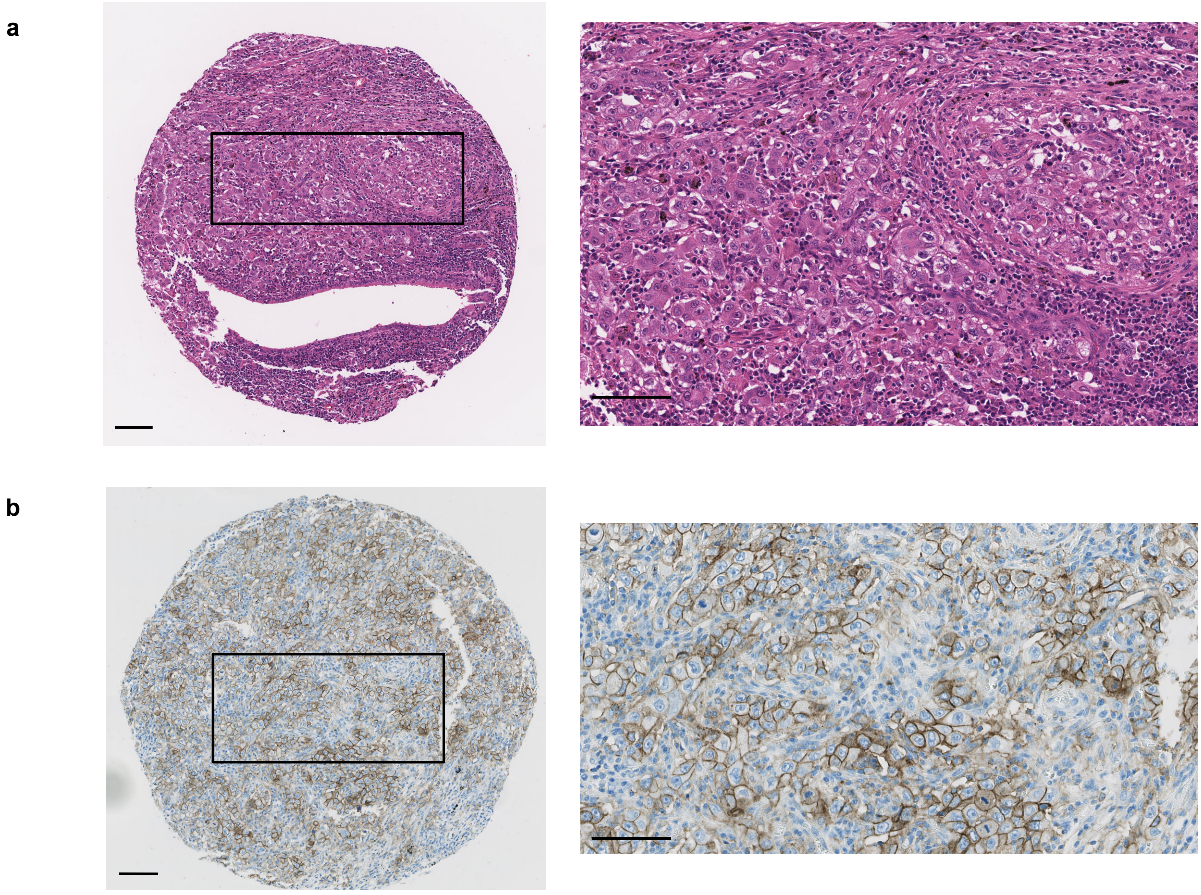

Supplementary Figure S7: (a) H&E-stained TMA core with high-risk hot discohesive morphology. Scale bars 100 $\mu$m. (b) PD-L1 immunohistochemistry of the same core. Scale bars 100 $\mu$m. PD-L1 programmed death ligand 1.

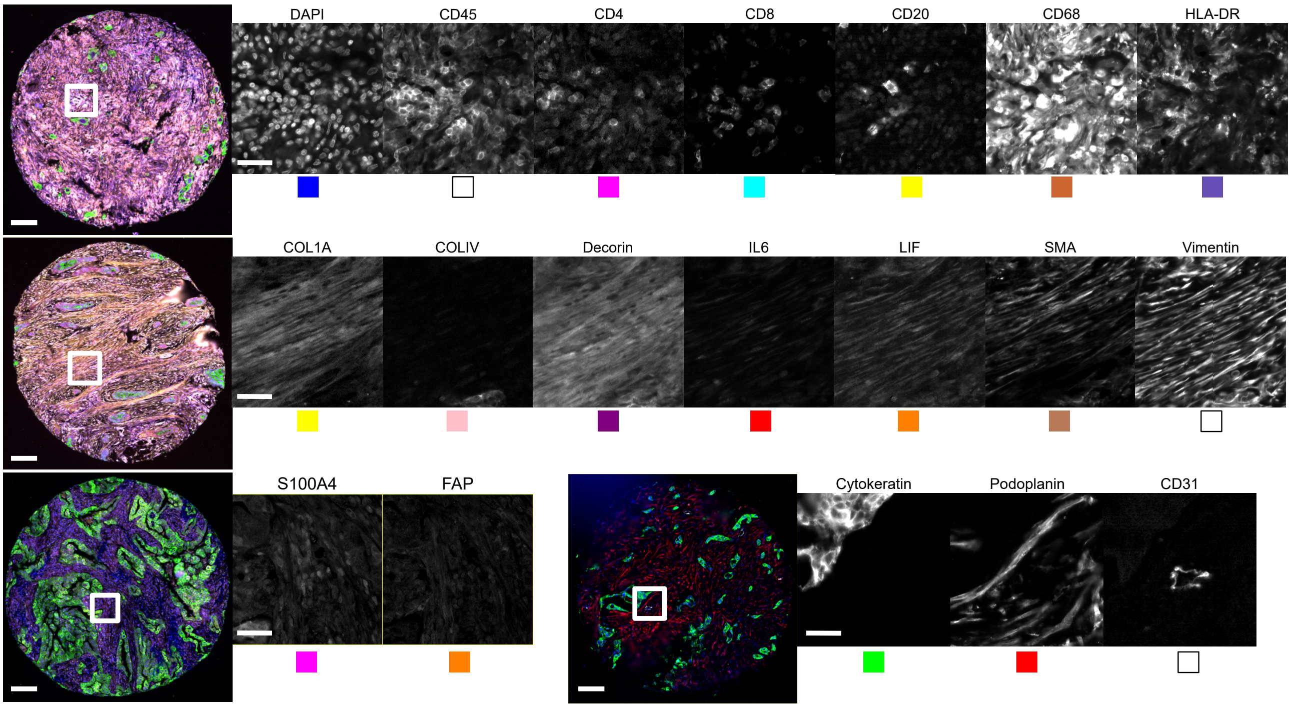

Supplementary Figure S8: Example mIF image from the fibroblast panel (see Figure 5.) Scale bars: core 100 $\mu$m, insert 30 $\mu$m. The coloured squares under the single channel image reference the colour on the merged image to the left of the single channel image.

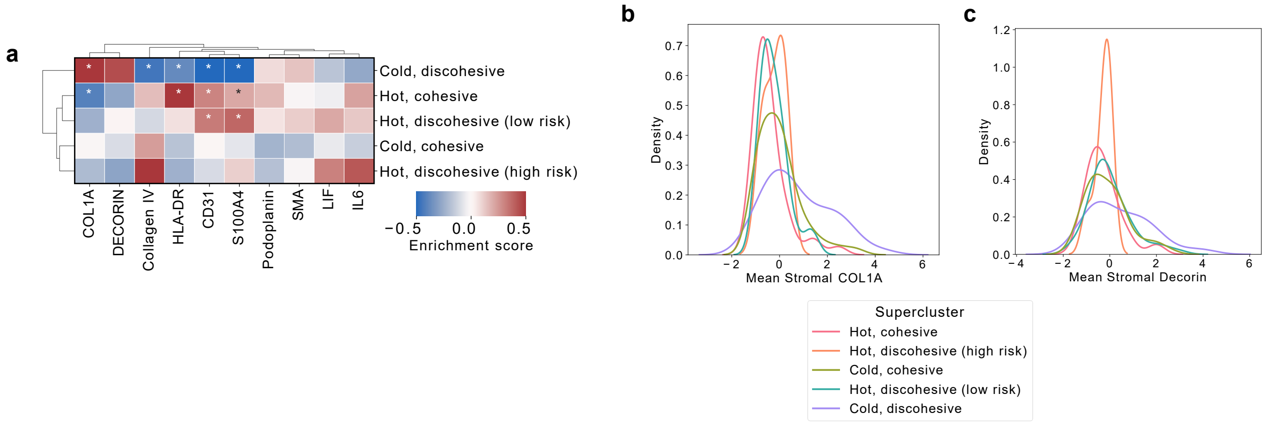

Supplementary Figure S9: (a) Enrichment of mean stromal intensity of stromal markers in the five superclusters; cold discohesive, n=44; hot cohesive, n=44; hot discohesive (low risk), n=50; cold cohesive, n=53; hot discohesive (high risk), n=7. Enrichment score calculated as the difference of means between the group and the population. Statistical significance tested with a two-tailed Mann-Whitney-U test and p-value correction for multiple testing using the Benjamini-Hochberg method (* $p < 0.05$). (b) Kernel density estimate plot showing the distribution of COL1A among the five superclusters. (c) Kernel density estimate plot showing the distribution of Decorin among the five superclusters.

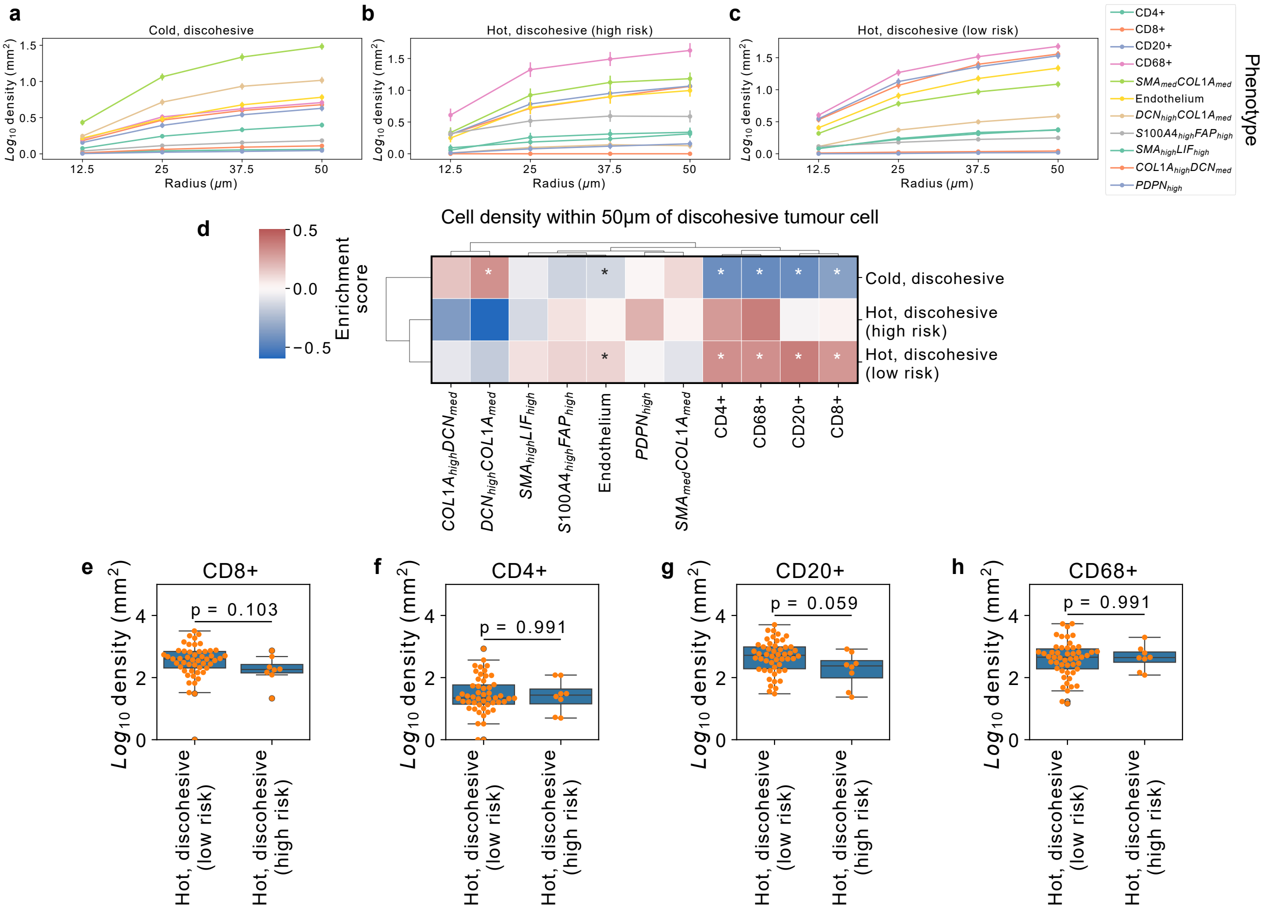

Supplementary Figure S10: (a-c) Mean density of stromal cells as a function of distance from discohesive tumour cells in the cold discohesive (a), low risk hot discohesive (b) and high risk discohesive (c) groups. Error bars are 95% confidence intervals. (d) Stromal cell density within 50 $\mu$m of a discohesive tumour cell; cold discohesive, n=44; hot discohesive (low risk), n=50; hot discohesive (high risk), n=8. Statistical significance tested with a two-tailed Mann-Whitney-U test and correction for multiple testing using the Benjamini-Hochberg method (* p $< 0.05$ (e-h) Density of CD8+ T-cells (e), CD4+ T-cells (f), CD20+ B-cells (g) and CD68+ macrophages (h) within 50 $\mu$m of a discohesive tumour cells in the two hot discohesive groups. Statistical significance tested with a two-tailed Mann-Whitney-U test.

Study **JLQ-001-24**
Experiment **JLQ-001-24_LUNG_ACA**
Starting well **A1**
Tissue type **Human FFPE**
Report date **9/3/2025 9:16:05 AM**

| Well | DAPI Channel | ATTO550 Channel | CY5 Channel | AF750 Channel |
|---|---|---|---|---|
| A1 | DAPI | CD105 - BX017 | -- - None | CD31 - BX001 |
| A2 | DAPI | FAP - BX002 | CD68 - BX015 | CD20 - BX007 |
| A3 | DAPI | CD38 - BX089 | CD45 - BX021 | -- - None |
| A4 | DAPI | Podoplanin - BX121 | HLA-DR - BX033 | GATA6 - BX028 |
| A5 | DAPI | -- - None | PDL1 - BX067 | COL1A - BX049 |
| A6 | DAPI | CXCL12 - BX041 | PD-1 - BX046 | CA9 - BX004 |
| A7 | DAPI | CD8 - BX026 | DECORIN - BX050 | SMA - BX013 |
| A8 | DAPI | E-cadherin - BX014 | CD4 - BX003 | -- - None |
| A9 | DAPI | PSTAT3 - BX029 | FOXP3 - BX031 | CXCL1 - BX034 |
| A10 | DAPI | FOXA1 - BX020 | Collagen IV - BX042 | GREM1 - BX025 |
| A11 | DAPI | VDR - BX023 | NCAD - BX027 | STAT3 - BX016 |
| A12 | DAPI | KI67 - BX047 | LIF - BX006 | Pan-Cytokeratin - BX019 |
| B1 | DAPI | S100A4 - BX052 | CTGF - BX054 | PDGFR - BX010 |
| B2 | DAPI | CRABP2 - BX055 | IL6 - BX036 | Vimentin - BX022 |
| B3 | DAPI | -- - None | HNF4A - BX045 | -- - None |
| B4 | DAPI | -- - None | CXCR4 - BX030 | -- - None |

**Blank cycles**

| Well | DAPI Channel | ATTO550 Channel | CY5 Channel | AF750 Channel |
|---|---|---|---|---|
| H1 | DAPI | -- - None | -- - None | -- - None |
| H2 | DAPI | -- - None | -- - None | -- - None |

Page 1 of 1

Supplementary Figure S11: Akoya PhenoCycler reporter template protocol

Supplementary Table 1: Patient Demographics

| **Characteristic** | **n** |
|---|---|
| Number of patients | 1007 |
| Number of images | 4427 |
| Median age at surgery (range) | 68 (31 - 89) |
| **Sex** | |
| Male | 468 (46.5%) |
| Female | 539 (53.5%) |
| **Tumour Stage** | |
| I | 359 (35.7%) |
| II | 198 (19.7%) |
| III | 278 (27.6%) |
| IV | 4 (0.4%) |
| Missing | 168 (16.7%) |
| **IASLC Grade** | |
| G1 | 169 (16.8%) |
| G2 | 294 (29.2%) |
| G3 | 544 (54.0%) |
| **2015 WHO Classification** | |
| Mucinous adenocarcinoma | 66 (6.6%) |
| Non-mucinous adenocarcinoma | 941 (93.4%) |

| Cluster | GP | Inflammation | Necrosis | S:E | Strom. cell. |
|---|---|---|---|---|---|
| 0 | Acinar | Mild-mod. | None | Equal | Moderate |
| 5 | Acinar | None-sparse | None | More stroma | Moderate |
| 11* | Acinar | Mild-mod. | None | More epithelium | Moderate |
| 13 | Solid | Mild-mod. | Some | More epithelium | Moderate |
| 17 | Solid | Mild-mod. | None | More epithelium | Moderate |
| 21 | Solid | Mild-mod. | None | Equal | Moderate |
| 27 | Acinar | None-sparse | None | More stroma | Moderate |
| 28 | Solid | Marked | None | Equal | Moderate |
| 30 | Solid | None-sparse | None | Equal | High |
| 31 | Acinar | None-sparse | None | Equal | Moderate |
| 33 | Acinar | Mild-mod. | Universal | Equal | Moderate |
| 34 | Cribriform | Mild-mod. | Some | More epithelium | Moderate |
| 35 | Lepidic | Mild-mod. | None | Equal | Moderate |
| 40 | Acinar | None-sparse | None | More stroma | Low |
| 41 | Papillary | Mild-mod. | None | Equal | Moderate |
| 44 | Acinar | Marked | None | Equal | High |
| 46 | Pap./Mpap. | None-sparse | None | More epithelium | Moderate |
| 47 | Solid | None-sparse | None | More epithelium | Moderate |
| 48 | Solid | Marked | Some | Equal | Moderate |
| 50* | Solid | Mild-mod. | None | Equal | Moderate |
| 52 | Lepidic | Mild-mod. | None | More epithelium | Moderate |
| 54 | Papillary | Mild-mod. | None | More epithelium | Moderate |
| 61 | Acinar | Marked | None | Equal | High |
| 64 | Acinar | Mild-mod. | None | More stroma | High |
| 67 | Acinar | Marked | None | Equal | Moderate |
| 68 | Acinar | Mild-mod. | None | Equal | Moderate |
| 69 | Acinar | Mild-mod. | None | More stroma | Moderate |

Supplementary Table 2: Clusters defined by malignant appearances, by consensus annotation. GP growth pattern; S:E stroma:epithelium ratio; Strom. cell. stromal cellularity. Pap. papillary; Mpap. micropapillary. * Clusters with less than high (>75%) purity

| Cluster | Feature | Inflammation | Necrosis |
|---|---|---|---|
| 1 | Collagenosis | None-sparse | None |
| 2 | Anthracosis | Mild-mod. | None |
| 3 | Collagenosis | None-sparse | None |
| 4 | Cartilage | None-sparse | None |
| 6 | Vessel | None-sparse | None |
| 7 | Necrosis | None-sparse | Some |
| 8 | Seromucinous glands | None-sparse | None |
| 9 | Necrosis | None-sparse | Universal |
| 10 | Lymphocytic stroma | Marked | None |
| 12 | Non-neoplastic pathological lung | None-sparse | None |
| 14 | Elastosis | None-sparse | None |
| 15* | Collagenosis | None-sparse | None |
| 16* | Lymphocytic stroma | Marked | Some |
| 18 | Elastosis | None-sparse | None |
| 19 | Normal/near-normal lung | Mild-mod. | None |
| 20 | Normal/near-normal lung | None-sparse | None |
| 22* | Vessel | None-sparse | None |
| 23* | Elastosis | None-sparse | Some |
| 24 | Necrosis | Mild-mod. | Universal |
| 25 | Lymphocytic stroma | Marked | None |
| 26 | Collagenosis | None-sparse | Some |
| 29 | Cartilage | None-sparse | None |
| 32 | Collagenosis | Marked | None |
| 36 | Normal/near-normal lung | None-sparse | None |
| 37 | Airways | None-sparse | None |
| 38 | Normal/near-normal lung | Mild-mod. | None |
| 39 | Normal/near-normal lung | Mild-mod. | None |
| 42 | Collagenosis | None-sparse | None |
| 43 | Collagenosis | None-sparse | None |
| 45 | Normal/near-normal lung | None-sparse | None |
| 49 | Vessel | Mild-mod. | None |
| 51 | Normal/near-normal lung | None-sparse | None |
| 53 | Normal/near-normal lung | None-sparse | None |
| 55 | Elastosis | None-sparse | None |
| 56 | Collagenosis | Mild-mod. | None |
| 57 | Normal/near-normal lung | None-sparse | None |
| 58 | Non-neoplastic pathological lung | Marked | None |
| 59 | Collagenosis | Mild-mod. | None |
| 60 | Collagenosis | Marked | None |
| 62 | Normal/near-normal lung | None-sparse | None |
| 63 | Normal/near-normal lung | Mild-mod. | None |
| 65 | Normal/near-normal lung | None-sparse | None |
| 66 | Normal/near-normal lung | None-sparse | None |
| 70 | Non-neoplastic pathological lung | Mild-mod. | None |

Supplementary Table 3: Remaining clusters, by consensus annotation. * Clusters with less than high (>75%) purity

| Primary Antibody | 1y Antibody Catalogue No. | Barcode | Barcode Catalogue No. |
|---|---|---|---|
| CD105/Endoglin (3A9) | (Cell Signaling #37995) | BX017 | Akoya 5250001 |
| Anti-Collagen I (EPR7785) | (Abcam ab215969) | BX042 | Akoya 4550122 |
| Anti-CRABP2 (EPR14256(B)) | (Abcam ab250468) | BX055 | Akoya 5250011 |
| Anti-CTGF (poly) | (Abcam ab6992) | BX054 | Akoya 5550019 |
| CXCL1 (poly) | (Proteintech 12335-1-AP) | BX034 | Akoya 5450007 |
| Anti-SDF1 (poly) | (Abcam ab9797) | BX041 | Akoya 5250008 |
| Decorin (E2N2C) | (Cell Signaling #55868) | BX050 | Akoya 5250016 |
| FAP (F1A4G) | (Abcam ab271976) | BX002 | Akoya 5450023 |
| Anti-FOXA1 (EPR10881-14) | (Abcam ab249749) | BX020 | Akoya 5250002 |
| GATA-6 (D61E4) | (Cell Signaling #48888) | BX028 | Akoya 5450005 |
| Human/Mouse Gremlin | (R&D Systems AF956) | BX025 | Akoya 5450004 |
| HNF4$\alpha$ (C11F12) | (Cell Signaling #31059) | BX045 | Akoya 5550016 |
| Anti-IL-6 antibody (1.2-2B11-2G10) | (Abcam ab9324) | BX036 | Akoya 5550014 |
| LIF Polyclonal antibody (poly) | (Proteintech 26757-1-ap) | BX006 | Akoya 5550018 |
| PDGF Receptor $\alpha$ (D1E1E) XP® | (Cell Signaling #32641) | BX010 | Akoya 5450021 |
| Phospho-Stat3 (Tyr705) | (Cell Signaling #9131) | BX029 | Akoya 5250005 |
| Anti-FSP1/S100A4 (poly) | (Sigma Aldrich 07-2274) | BX052 | Akoya 5250012 |
| Stat3 (124H6) | (Cell Signaling #83541) | BX029 | Akoya 5450001 |
| VDR Monoclonal (9A7) | (Invitrogen MA1-710) | BX023 | Akoya 5250003 |
| Anti-Carbonic Anhydrase 9/CA9 (poly) | (Abcam ab15086) | BX004 | Akoya 5450019 |
| Anti-N Cadherin (EPR1791-4) | (Abcam ab271856) | BX027 | Akoya 5550011 |
| Pre-Conjugated Antibodies | | | |
| Anti-Hu CD4 (AKYP0048)—Alexa Fluor™ 647 | (Akoya Biosciences 4550112) | BX003 | N/A |
| Anti-Hu CD8(AKYP0028)—Atto 550 | Akoya Biosciences 4250012) | BX026 | N/A |
| Anti-Hu CD20(AKYP0049)—Alexa Fluor™ 750 | (Akoya Biosciences 4450094) | BX064 | N/A |
| Anti-Hu CD31(AKYP0047)—Alexa Fluor™ 750 | (Akoya Biosciences 4450017) | BX001 | N/A |
| Anti-Hu CD38(AKYP0110)—Atto 550 | (Akoya Biosciences 4250080) | BX089 | N/A |
| Anti-Hu CD45(AKYP0074)—Alexa Fluor™ 647 | (Akoya Biosciences 4550121) | BX021 | N/A |
| Anti-Hu CD68(AKYP0050)—Alexa Fluor™ 647 | (Akoya Biosciences 4550113) | BX015 | N/A |
| Anti-Hu FOXP3(AKYP0102)—Alexa Fluor™ 647 | (Akoya Biosciences 4550071) | BX031 | N/A |
| Anti-Hu HLA-DR(AKYP0063)—Alexa Fluor™ 647 | (Akoya Biosciences 4550118) | BX033 | N/A |
| Anti-Hu/Mu Ki67(AKYP0052)—Atto 550 | (Akoya Biosciences 4250019) | BX047 | N/A |
| Anti-Hu Pan-Cytokeratin(AKYP0053)—Alexa Fluor™ 750 | (Akoya Biosciences 4450093) | BX066 | N/A |
| Anti-Hu PD-1(AKYP0070)—Alexa Fluor™ 647 | (Akoya Biosciences 4550038) | BX046 | N/A |
| Anti-Hu PD-L1(AKYP0103)—Alexa Fluor™ 647 | (Akoya Biosciences 4550128) | BX067 | N/A |
| Anti-Hu Podoplanin (AKYP0007)—Atto 550 | (Akoya Biosciences 4250094) | BX121 | N/A |
| Anti-Hu SMA(AKYP0081)—Alexa Fluor™ 750 | (Akoya Biosciences 4450049) | BX013 | N/A |
| Anti-Hu Vimentin(AKYP0082)—Alexa Fluor™ 750 | (Akoya Biosciences 4450050) | BX022 | N/A |
| Anti-Hu Collagen IV(AKYP0083)—Alexa Fluor™ 647 | (Akoya Biosciences 4550122) | BX042 | N/A |
| Anti-Hu E-cadherin (AKYP0057)—Atto 550 | (Akoya Biosciences 4250021) | BX014 | N/A |

Supplementary Table 4: LATTICeA fibroblast panel antibodies and barcodes

| Conjugation |
|---|
| 1. Add 500 $\mu$l of Filter blocking solution (Antibody Conjugation Kit: Akoya Biosciences, Ref. 7000009) to the top of the 50kDa MWCO filter (Sigma Aldrich, Ref. UFC5050) and spin down at 12000g for 2 mins.<br>2. Empty both the top and the bottom of the column.<br>3. Add 100 $\mu$g of the antibody and 300 $\mu$l of PBS.<br>4. Spin down at 12000g for 8 mins empty the bottom of the column.<br>5. Create the antibody reduction master mix: Reduction solution 2: 275 $\mu$l + Reduction solution 1: 6.6 $\mu$l (Antibody Conjugation Kit: Akoya Biosciences, Ref. 7000009).<br>6. Add 260 $\mu$l of the reduction master mix, pipette up and down or vortex for 3 sec.<br>7. Incubate for 15 minutes at RT.<br>8. Add 200 $\mu$l of conjugation solution (Antibody Conjugation Kit: Akoya Biosciences, Ref. 7000009).<br>9. Spin down at 12000g for 8mins empty the bottom of the column.<br>10. Add 450 $\mu$l of conjugation solution pipette up and down or vortex for 3 sec.<br>11. Spin down at 12000g for 8mins empty the bottom of the column<br>12. Prepare the barcode solution:<br>13. Resuspend two barcodes, each in 10 $\mu$l of nuclear free water (Thermo Scientific, Ref. R0582). Avoid vortex.<br>14. Add 210 $\mu$l of Conjugation solution to each suspended Barcode (avoid vortex, gentle pipette)<br>15. Leave on the shaker<br>16. Add the barcode solution to the top of the column that contain the antibody, pipette up and down or vortex for 3 sec.<br>17. Transfer in a low bind tube and place on the shaker and incubate overnight at 4℃<br>18. Transfer the solution back in the column<br>19. Spin down at 12000g for 8 mins empty the bottom of the column<br>20. Add 450 $\mu$l of purification solution (Antibody Conjugation Kit: Akoya Biosciences, Ref. 7000009), pipette up and down or vortex for 3 sec.<br>21. Spin down at 12000g for 8 mins empty the bottom of the column<br>22. Add 450 $\mu$l of purification solution, pipette up and down or vortex for 3 sec.<br>23. Spin down at 12000g for 8 mins empty the bottom of the column<br>24. Add 450 $\mu$l of purification solution, pipette up and down or vortex for 3 sec.<br>25. Spin down at 12000g for 8 mins empty the bottom of the column<br>26. The top of the column will contain the purified solution.<br>27. Add 150 $\mu$l of Antibody Stabilizer PBS PROTEIN-FREE (Bioaxxess, Ref. 331), pipette up and down or vortex for 3 sec.<br>28. Place a new empty tube upside-down on top of the filter unit column.<br>29. Invert the filter for collection into the new collection tube<br>30. Spin down at 3000g for 2 mins.<br>31. The final volume should be around 170 $\mu$l.<br>32. Transfer in an Eppendorf.<br>33. Conjugated antibodies are validated in both DAB and fluorescence to ensure specificity and sensitvity.<br>Staining and Image Generation<br>34. Manual staining according to the Akoya Phenocycler-Fusion user manual.<br>35. The first antibody cocktail was applied for 3 hours at room temperature with the following antibody-barcode pairings (Supplementary Table 6).<br>36. The second cocktail was applied overnight at 4℃ with the following antibody-barcode pairings (Supplementary Table 6).<br>37. Akoya Reporters were applied following the protocol template (Supplementary Figure S11), with an adjustment to the Nuclear Stain (10 $\mu$l per well at 5ms). Exposure times were kept at 150ms for each marker and experiment was run at 8-bit. |

38. Images were QC'd using QuPath Version 0.5.0.

Supplementary Table 5: Modified Phenocycler Fusion protocol used for the LATTICeA fibroblast panel.

| Primary Antibody | Antibody-Barcode Dilution |
|---|---|
| First primary antibody-barcode cocktail incubation | |
| CD105/Endoglin (3A9) - BX017 | 1:10 |
| Anti-Hu CD31 - BX001 | 1:200 |
| Anti-N Cadherin antibody - BX027 | 1:50 |
| Anti-Hu CD20 - BX007 | 1:500 |
| Anti-Hu CD38 - BX089 | 1:200 |
| Anti-Hu CD45 - BX021 | 1:300 |
| Anti-Hu E-cadherin - BX014 | 1:200 |
| Anti-Hu Podoplanin - BX121 | 1:100 |
| Anti-Hu HLA-DR - BX033 | 1:300 |
| Anti-Carbonic Anhydrase 9/CA9 - BX004 | 1:25 |
| Anti-Hu PD-L1 - BX067 | 1:50 |
| Anti-SDF1 antibody - BX041 | 1:50 |
| Anti-Hu PD-1 - BX046 | 1:50 |
| Anti-Hu Collagen IV - BX042 | 1:200 |
| Anti-Hu CD8 - BX026 | 1:100 |
| Decorin (E2N2C) XP® - BX050 | 1:100 |
| Anti-Hu SMA - BX013 | 1:300 |
| Anti-Hu CD4 - BX003 | 1:100 |
| Anti-Hu FOXP3 - BX031 | 1:50 |
| CXCL1 - BX034 | 1:100 |
| Anti-FOXA1 - BX020 | 1:1000 |
| VDR - BX023 | 1:200 |
| Anti-Hu/Mu Ki67 - BX047 | 1:100 |
| LIF - BX006 | 1:200 |
| Anti-Hu Pan-Cytokeratin - BX066 | 1:600 |
| Anti-CTGF - BX054 | 1:300 |
| Anti-Hu Vimentin - BX022 | 1:300 |
| Anti-CRABP2 - BX055 | 1:25 |
| Anti-IL-6 - BX036 | 1:200 |
| Anti-Hu CD68 - BX015 | 1:800 |
| Second overnight primary antibody-barcode cocktail incubation | |
| Anti Collagen I - BX049 | 1:25 |
| FAP - BX002 | 1:10 |
| GATA6 - BX028 | 1:10 |
| Gremlin - BX025 | 1:25 |
| HNF4$\alpha$ - BX045 | 1:100 |
| PDGF Receptor $\alpha$ - BX010 | 1:25 |
| Phospho STAT3 - BX029 | 1:10 |
| Anti FSP1/S100A - BX052 | 1:25 |
| STAT3 - BX016 | 1:50 |
| Anti CXCR4 - BX030 | 1:25 |

Supplementary Table 6: Primary antibody-barcode cocktail incubations (LATTICeA fibroblast panel)
